# A Spatially Structured Spiking Network Model of Beta Traveling Waves and Their Attenuation in Motor Cortex

**DOI:** 10.64898/2026.03.18.712701

**Authors:** Ludovica Bachschmid-Romano, Nicholas Hatsopoulos, Nicolas Brunel

**Affiliations:** Department of Neurobiology, Duke University, Durham, NC, USA; INMED, Aix Marseille University, Marseille, France; Department of Anatomy and Organismal Biology, University of Chicago, Chicago, IL, USA; Department of Physics, Duke University, Durham, NC, USA; Department of Computing Sciences, Bocconi University, Milan, Italy

**Author notes:** Co-senior authors (Nicholas Hatsopoulos, Nicolas Brunel).

## Abstract

Beta-band oscillations in primate motor cortex propagate as planar traveling waves whose amplitude attenuates with spatial gradients across the cortical sheet just before movement onset. How local excitatory–inhibitory (E–I) interactions and spatial connectivity jointly generate these waves, their attenuation patterns, and their stereotyped rostro–caudal bias remains unclear. Here we address this question by implementing a spatially structured network of leaky integrate-and-fire neurons with distance-dependent connectivity, conduction delays, and realistic synaptic dynamics. Through linear stability analysis and large-scale simulations validated against macaque electrophysiology, we show that planar beta waves emerge as Turing–Hopf spatiotemporal instabilities, where global beta oscillations coexist with irregular single-neuron firing. When the network operates near the boundary between oscillatory and asynchronous regimes, internally generated fluctuations produce the irregular, transient beta bursts characteristic of single-trial local field potentials. A rapid, spatially homogeneous increase in external drive pushes the circuit into an asynchronous state, reproducing the beta power reduction and spatial attenuation gradients seen at movement onset, alongside the irregular spatiotemporal dynamics of movement execution. By introducing anisotropic excitatory-to-excitatory connectivity, we recover the observed rostro–caudal propagation bias. Our results suggest that motor cortical traveling waves are intrinsic dynamical modes of local E–I circuits, recruited and modulated by behaviorally relevant inputs to organize movement initiation.

## 1 Introduction

Large-scale propagating activity patterns are a widespread feature of cortical dynamics across species and behavioral contexts [1, 2]. In primate motor cortex, local field potentials (LFPs) recorded from multielectrode arrays reveal transient beta-band (15–30 Hz) oscillations that propagate as planar waves along the rostro–caudal axis [3]. Single-trial analysis [4] showed that beta power attenuates immediately prior to movement onset, and the timing of beta attenuation forms a spatial gradient across the cortical sheet. These gradients are time-locked to movement initiation, suggesting that propagating beta attenuation reflects evolving cortical excitability preceding voluntary movement.

Although beta oscillations in motor cortex have traditionally been attributed either to inputs from basal ganglia–thalamic loops [5, 6] or to intrinsic cortical circuitry [7, 8], recent work points to an intermediate viewpoint: transient beta events arise locally from the interaction of excitatory and inhibitory microcircuits, but are initiated and shaped by excitatory inputs — likely relayed via thalamic pathways influenced by basal ganglia [9]. An open question is how these local microcircuit mechanisms interact with the large-scale spatial structure of cortical connectivity to generate organized planar waves. Previous modeling studies have laid the groundwork to address these questions. Neural-field and rate models have shown that spatially structured lateral excitation and inhibition can destabilize homogeneous activity and generate pattern-forming instabilities, including traveling and standing waves [10, 11, 12, 13, 14, 15, 16, 17]. Subsequent analyses incorporating finite conduction delays have derived dispersion relations that characterize Turing, Hopf, and mixed Turing–Hopf instabilities and the resulting spatiotemporal regimes [18, 19, 20, 21]. Related spatiotemporal instabilities have also been demonstrated in spatially extended balanced spiking networks, where distance-dependent connectivity and the relative spatial spread of excitatory and inhibitory projections control the emergence of patterned activity [22, 23, 24]. Balanced spiking networks with distance-dependent connectivity have also been shown to generate irregular propagating activity consistent with the low-rate, weakly correlated asynchronous–irregular dynamics observed in cortex, while reproducing the spatiotemporal statistics of marmoset area MT [25, 26]. More recently, a rate model with locally oscillatory excitatory–inhibitory modules coupled by long-range excitation was used to show that beta bursts and traveling waves in primate motor cortex can arise from the spatial dephasing of an otherwise synchronized beta rhythm by structured stochastic external inputs [27].

Yet it remains unanswered whether a network with realistic connectivity can generate planar waves during movement preparation, attenuate them prior to movement onset, produce spatial gradients in beta-attenuation timing, and exhibit the experimentally observed propagation bias, while also reproducing the spatiotemporal statistics of multi-unit activity during both movement preparation and execution.

Here, we address this question using a spatially structured network of leaky integrate-and-fire neurons with distance-dependent connectivity, conduction delays, and realistic AMPA and GABA synaptic kinetics. This architecture provides the excitatory–inhibitory interaction timescales needed for beta-band oscillations [28, 29] and enables a direct link between circuit structure and LFP-based measurements. Through linear stability analysis of the asynchronous state and large-scale simulations, we show that planar beta waves emerge as Turing–Hopf spatiotemporal instabilities at intermediate levels of external drive, where global beta oscillations coexist with irregular single-neuron firing. The model reproduces experimental data from macaque motor cortex when operating close to the boundary between oscillatory and asynchronous regimes, so that internally generated fluctuations determine when the unstable beta mode is expressed. This is consistent with experimental reports that beta activity appears as transient bursts [9], and with previous modeling work showing that proximity to a bifurcation can give rise to intermittent beta activity [27]. We find that increasing external input – representing the transition from motor preparation to execution – drives the circuit through a bifurcation into an asynchronous state, reproducing the beta power attenuation observed at movement initiation, with characteristic spatial gradients in beta attenuation times. Across these regimes, the model recapitulates the spatiotemporal statistics of LFP activity observed during both movement planning and execution. Finally, we show that anisotropic recurrent connectivity gives rise to directional wave propagation, without requiring spatially heterogeneous external inputs. By selectively introducing anisotropy in different synaptic pathways, we identify anisotropic excitatory-to-excitatory connectivity as a parsimonious and testable mechanism underlying the rostro–caudal propagation bias observed in motor cortex.

## 2 Results

To investigate the mechanistic origin of cortical traveling waves, we consider a two-population excitatory–inhibitory network distributed over a two-dimensional cortical sheet, with distance-dependent connectivity and synaptic delays. The network has a sparse recurrent synaptic connectivity, with a connection probability that decreases exponentially with the distance between neurons over a range of a few hundred micrometers. We analyze and simulate networks with both isotropic and anisotropic spatial connectivity profiles. The spatial spread of *E* → *E* connections is broader than that of *E* → *I, I* → *E*, and *I* → *I* projections, consistent with anatomical and physiological studies in mammalian cortex [30, 31, 30, 32]. Synaptic currents have finite rise and decay times, described as delayed difference of exponential kernels with distinct AMPA and GABA time constants. Consistent with cortical physiology, AMPA currents decay faster than GABAergic currents [33, 34, 35], while the membrane integration time constant of excitatory neurons is longer than that of fast-spiking interneurons [36, 37]. This establishes the excitatory–inhibitory interaction timescale needed to generate beta-band oscillations. The characteristics of spatial connectivity and synaptic timescales are compatible with theoretical results showing that networks of LIF neurons exhibit stable asynchronous, irregular activity, similar to that observed in cortical recordings, when excitation extends over broader spatial ranges than inhibition [20, 22, 23] and inhibitory synapses are slower than excitatory ones [38]. All neurons receive additional stochastic external inputs modeled as spatially homogeneous Poisson spike trains, mimicking inputs from thalamic and cortico-cortical projections. To model LFPs, we draw on results from detailed biophysical modeling studies [39], which compared LFPs generated by morphologically detailed multi-compartment neuron models with those estimated from point-neuron LIF networks. These studies showed that a weighted sum of the absolute values of AMPA and GABA currents onto excitatory neurons provides an accurate and computationally efficient approximation of the LFP. For layer 2/3 pyramidal cells, the contribution of synaptic currents from neurons within a radius of *r* < *r*_x_ ~ 100*µ*m was estimated to slowly decay with the distance to the electrode, while the contribution of neurons placed at larger distances *r* > *r*_x_ decays as ~ 1*/r*^2^ [40]. A schematic of the model is provided in Fig. 1.a, and details of the model and of LFP proxies are given in Sections 4.1 and 4.2; parameter values are summarized in Table 1.

**Table 1:** Neuron and synapse parameter values.

| Membrane & Refractory |  |
| --- | --- |
| $V_\theta$ | 18 mV |
| $V_{\text{reset}}$ | 11 mV |
| $\tau_m^E$ | 20 ms |
| $\tau_m^I$ | 10 ms |
| $\tau_{\text{rp}}^E$ | 2 ms |
| $\tau_{\text{rp}}^I$ | 1 ms |
| Delays |  |
| $\tau_0$ | 0.1 ms |
| $v$ | 0.3 mm/ms |
| $\bar{\tau}_L^{EE}$ | 2.57 ms |
| $\bar{\tau}_L^{EI}, \bar{\tau}_L^{IE}, \bar{\tau}_L^{II}$ | 1.725 ms |
| Synaptic time constants |  |
| $\tau_{\text{ref}}^E$ | 2 ms |
| $\tau_{\text{ref}}^I$ | 1 ms |
| $\tau_r^{AE}$ | 0.7 ms |
| $\tau_d^{AE}$ | 3.5 ms |
| $\tau_r^{AI}$ | 0.7 ms |
| $\tau_d^{AI}$ | 3.5 ms |
| $\tau_r^G$ | 1 ms |
| $\tau_d^G$ | 18 ms |
| Synaptic weights |  |
| $J_{\text{ext}}^I$ | 0.85 mV |
| $J_{\text{ext}}^E$ | 0.455 mV |
| $g$ | 3.6 |
| $w_{EE}$ | 0.1575 mV |
| $w_{IE}$ | 0.2625 mV |
| $w_{EI}$ | $1.07 g w_{EE}$ |
| $w_{II}$ | $g w_{IE}$ |
| Peak amplitude of EPSP and IPSP of the membrane potential at g=4.1 |  |
| $\tilde{J}_{EE}$ | 0.1084 mV |
| $\tilde{J}_{IE}$ | 0.1480 mV |
| $\tilde{J}_{EI}$ | 0.5106 mV |
| $\tilde{J}_{II}$ | 0.6244 mV |
| Connectivity |  |
| $C_{EE}$ | 800 |
| $C_{IE}$ | 800 |
| $C_{EI}$ | 200 |
| $C_{II}$ | 200 |
| $l_{EE}$ | 380 $\mu\text{m}$ |
| $l_{IE} = l_{EI} = l_{II}$ | 250 $\mu\text{m}$ |
| $\rho_{EE}$ | 0.6 |
| $\rho_{IE} = \rho_{EI} = \rho_{II}$ | 0 |
| External input in Fig 1-2 |  |
| $\sigma_n$ | 3.3 Hz/$\sqrt{\text{ms}}$ |
| $\tau_n$ | 150 ms |

**Figure 1:**
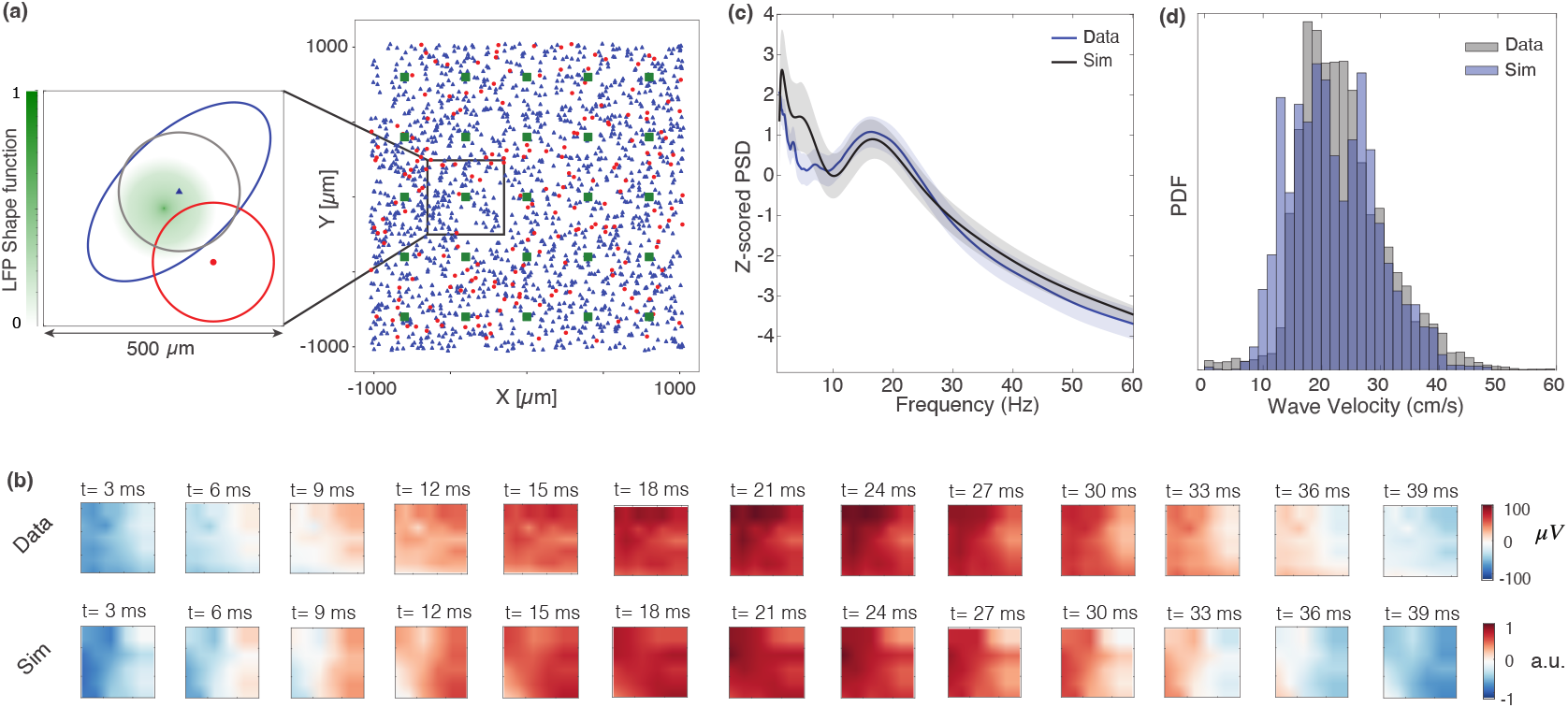
Comparison of LFP activity from recordings and simulations. (a) Schematic of the model. Right: a random subset of neurons shown at their locations on the 2D sheet. Blue triangles correspond to excitatory neurons, red dots to inhibitory neurons. Green squares mark virtual recording sites, where LFP proxies are computed (25 sites arranged in a 5 × 5 grid). Left: spatial spread of connection probabilities for a model with anisotropic E to E connectivity – blue ellipse for E→E, gray circle for E→I, and red circle for both I→I and I→E (see Table 1 for parameters). The green shade shows the spatial profile of the LFP shape function. (b) Top: example of LFPs in the beta band propagating along the diagonal electrodes of the array as planar traveling wave. For the purpose of comparison with simulations, the top row shows a section of the Utah array with 5×5 channels. Bottom: example of a traveling wave from simulations, with virtual recording sites placed every 400 *µ*m as shown in panel (a). (c) Normalized power spectral density, mean ± standard deviation over 30 trials, from recordings and simulations. The empirical PSD (black) was smoothed by convolution with a Gaussian kernel (FWHM of ~ 2*Hz* and ±3*σ* support). (d) Velocities of detected plane waves from recordings and simulations.

### Beta waves: model vs experiment

With appropriately chosen parameters, our model reproduces salient features of multielectrode recordings from the primary motor cortex of monkeys during an instructed-delay reaching task [3]. For a specific range of external input strengths, the population displays prominent beta-band oscillations that spontaneously propagate in space (Fig. 1b, bottom), closely reproducing the planar traveling waves recorded from motor cortex when the animal is at rest (Fig. 1b, top). Fig. 1c shows that the model captures the main spectral features of the experimental data, with both power spectral densities exhibiting a clear peak within the same beta-frequency range. Finally, Fig. 1d demonstrates that the distribution of detected wave velocities in the model is comparable to that observed experimentally.

### Spatiotemporal patterns of beta attenuation at movement onset

We next turn to how beta waves depend on inputs to the network. Balasubramanian et al. [4] showed that beta-band activity is high during rest and movement preparation but attenuates sharply just before movement onset (Fig. 2-a). The timing of this attenuation is locked to movement initiation rather than to the Go cue, indicating a mesoscopic signature of movement initiation. Moreover, the attenuation does not occur simultaneously across the cortical sheet but forms spatial patterns that propagate a few hundred milliseconds before movement begins (Fig. 2-c,d). In our model, these spatial patterns of beta attenuation can emerge from a rapidly increasing, spatially homogeneous external input. As the strength of this external drive increases (Fig. 2b), the network transitions from a beta-range oscillatory regime to a more asynchronous state, leading to a corresponding drop in beta power. On a single-trial basis, spatial gradients of beta attenuation emerge at the conclusion of the final beta wave (Fig. 2e,f). These gradients align with the direction of the wave gradient, and closely mirror the attenuation patterns observed in motor cortical recordings prior to movement. Fig. 2g,h shows examples of LFP amplitude envelopes from single trials in both recordings and simulations, alongside their corresponding autocorrelation functions (ACs). The ACs calculated over the rest period (yellow area, corresponding to lower external input in the model) exhibit clear, damped oscillations in the beta band. Following movement onset (blue area, induced by an increase in external input in the model), this oscillatory structure significantly dissipates, reflecting a transition to a more aperiodic regime in both the biological and simulated networks.

**Figure 2:**
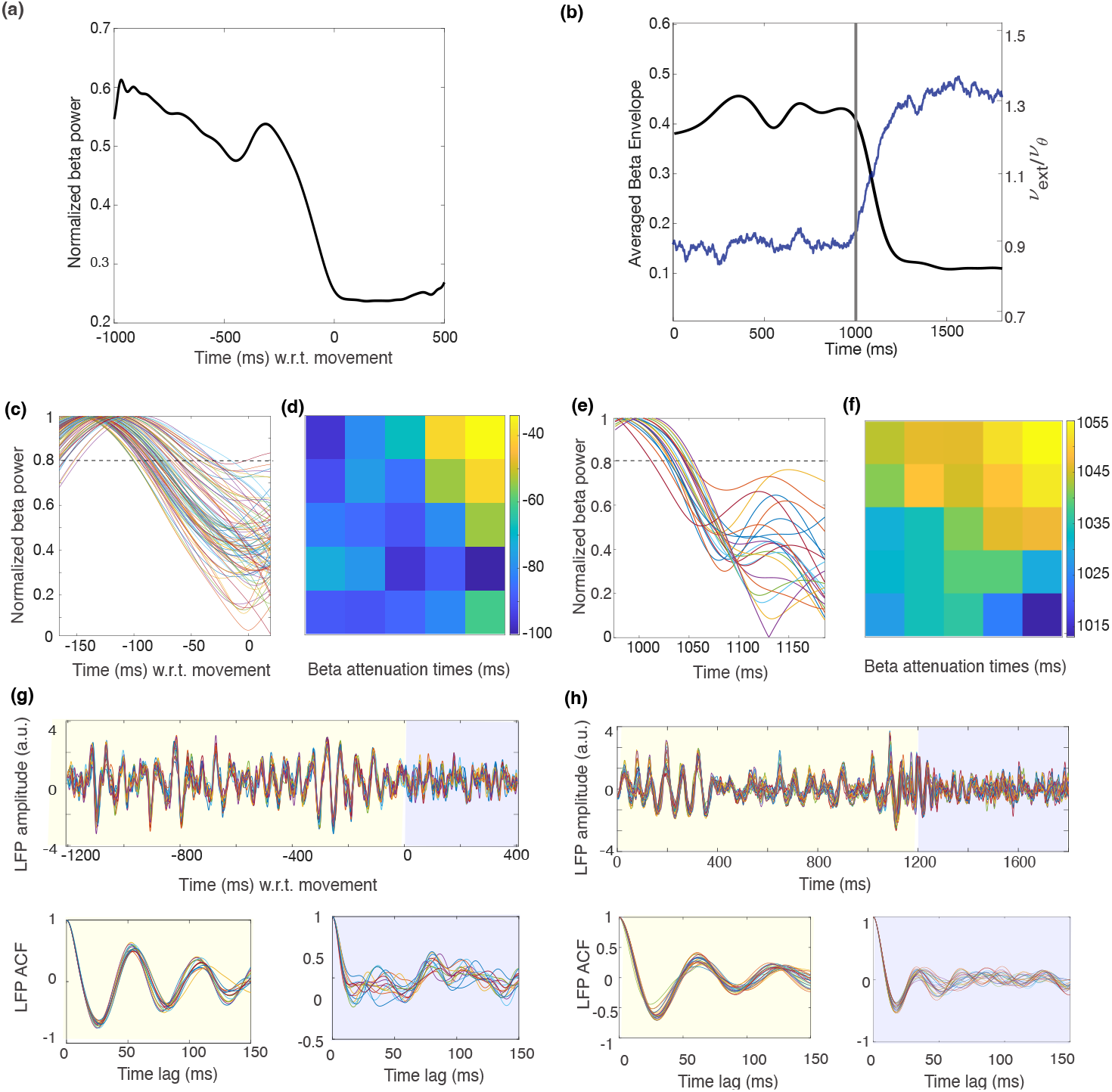
Spatio-temporal patterns of beta attenuation. (a) Temporal profile of beta power averaged over 200 trials aligned on movement onset, from recordings. (b) Temporal profile of beta power from simulations, averaged over 30 trials (black line), and temporal evolution of the external population rate, expressed relative to the threshold rate *ν*_θ_ required to reach threshold in the absence of recurrent input (blue line; scale shown on the right axis). (c) Single-trial LFP normalized beta envelopes from recordings (all channels), as a function of time since movement onset. The beta attenuation time is defined as the point where the envelope crosses the threshold (dashed line). (d) Spatial map of beta attenuation timings on the array. For comparison with simulations, the panel shows a section of the Utah array with 5 × 5 channels. (e-f) As in (c-d), but for the activity from simulations. (e) Band-pass filtered LFP amplitude in the [10,80] Hz range for one trial of the center-out reaching task, from cue onset to the end of movement (top). Autocorrelation of the LFP for the same trial during movement preparation – defined as the interval before movement onset – and during movement execution (bottom). (f) Same quantities as in (e), but computed from simulations using the time-varying external input shown in panel (b). Lower input levels yield traces resembling movement preparation, whereas higher input levels resemble movement execution. In simulations, movement onset is defined as the time at which the average beta power reaches its baseline level (panel b), in analogy with the experimental data (panel a).

Overall, the model demonstrates that beta waves and spatiotemporal patterns of beta attenuation can emerge intrinsically from the local circuit dynamics and connectivity structure, without requiring spatially patterned external inputs.

### Analysis of the model: Dynamical regimes

For different parameter values, the model exhibits a range of spatiotemporal regimes. To delineate the domains of asynchronous–irregular activity and synchronous–irregular activity (i.e., global oscillations with irregular spiking), we perform a linear stability analysis following the framework of [41, 42, 43]. We consider the mean-field limit on an infinite continuous sheet and compute the equilibrium firing rates 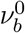 for excitatory (*b* = *E*) and inhibitory (*b* = *I*) populations. By introducing a small plane-wave perturbation of the presynaptic rate,

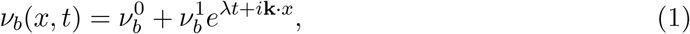

where *λ* = *α* + *iω*, we derive the resulting modulations in the mean and variance of the postsynaptic input currents. These input fluctuations are converted into postsynaptic firing rate perturbations 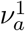 via single-neuron susceptibilities to mean and variance modulations (see (28)). Self-consistency requires that the postsynaptic perturbations match the imposed presynaptic ones, leading to the 2×2 eigenvalue problem ***ν***^1^ = 𝒜(*λ*, **k**)***ν***^1^ (where *A* is defined in (29)). The stability of the system is thus governed by the characteristic equation:

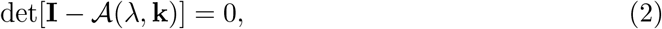

which implicitly defines the dispersion relation *λ*(**k**) = *α*(**k**) + *iω*(**k**). Because the synaptic filters in our model have finite rise and decay times, the input drive in the mean-field limit is colored with a finite correlation time. To account for this, we utilized numerical susceptibilities and a parametric mapping function to correct the analytical white-noise limits. Detailed derivations and plots of these colored-noise susceptibilities (Fig. S1) are provided in the SI Appendix.

We solved numerically (2) to characterize the model dynamical regimes, by outlining the region of stability of asynchronous state, and computing the spatio-temporal characteristics of the instabilities of this state. Fig. 3 reports results for a spatially homogeneous system in which connectivity and delays are independent of position, and perturbations are purely temporal (**k** = **0**). Connection probabilities are chosen to match the average indegree of the full model, and delays are set to the corresponding spatially averaged effective delays of the full model. Fig. 3.a shows the phase diagram as a function of the external input strength *ν*_ext_ and the inhibitory gain parameter *g*, which scales inhibitory synaptic strengths (larger *g* strengthens GABAergic input to both populations). For the range of *g* shown, low *ν*_ext_ yields a low-rate asynchronous-irregular (AI) state (Fig. 3c, top). In the intermediate band (*g* ≲ 4.2), increasing *ν*_ext_ triggers a Hopf bifurcation into global oscillations (Fig. 3c, middle), though single-neuron spiking remains irregular. At higher *ν*_ext_, a second Hopf bifurcation occurs, beyond which the system returns to a high-rate AI state (Fig. 3c, bottom). Across both regimes, the inter-spike interval coefficient of variation (CV) remains near unity (Fig. S2), confirming that single-neuron firing remains irregular regardless of global synchrony. Stationary firing rates from mean-field theory show close agreement with simulations (Fig. 3b, top). Similarly, the beta-range oscillation frequency predicted by linear stability analysis matches that measured in simulations (Fig. 3b, bottom).

**Figure 3:**
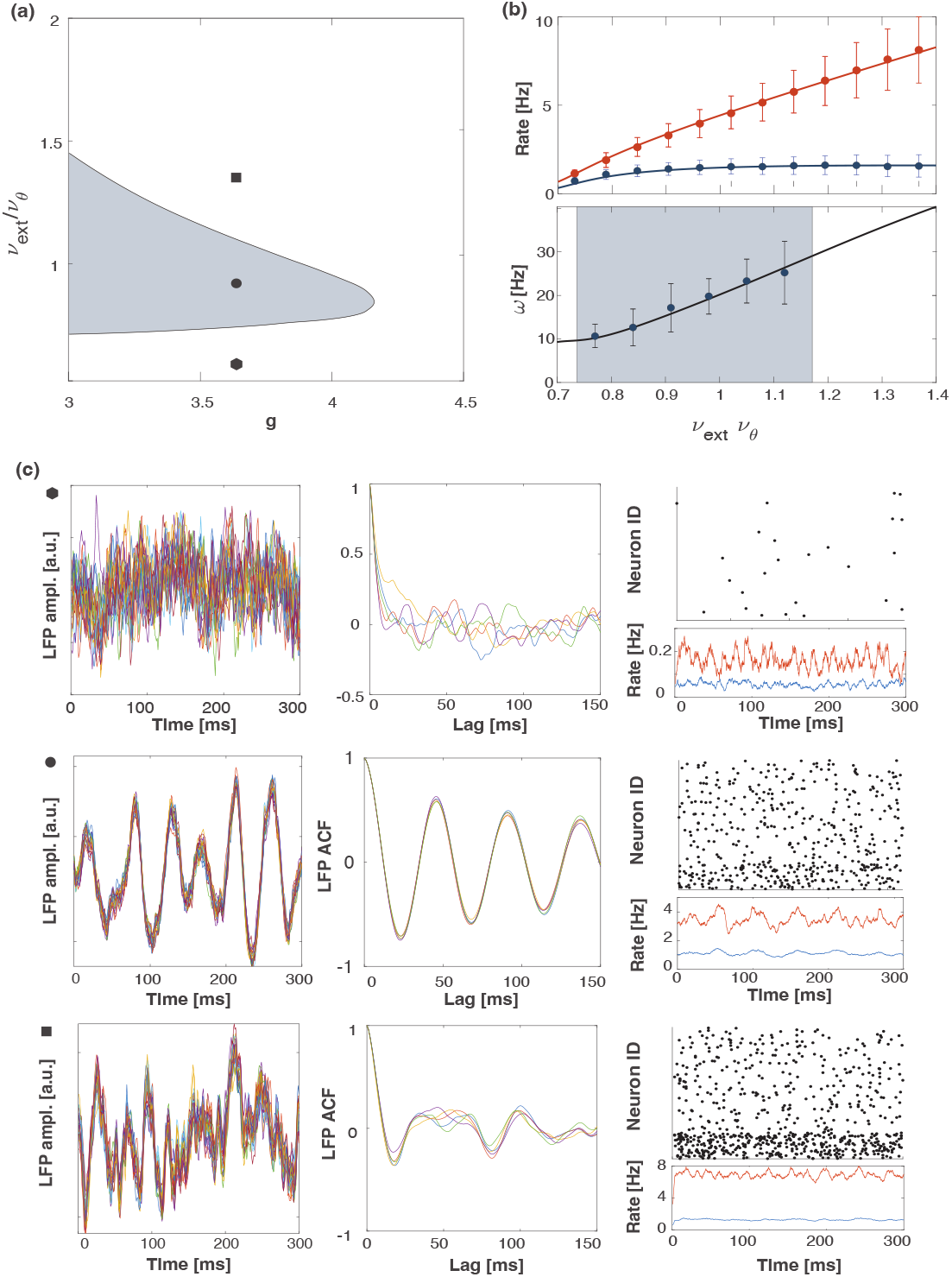
Analysis of the temporal dynamics in a spatially homogeneous model. (a) Bifurcation diagram for the spatially homogeneous network as a function of the external input strength *ν*_*ext*_*/ν*_θ_ and the excitation-inhibition ratio *g*. Larger *g* strengthens inhibitory inputs to both E and I populations. Gray area: synchronous irregular activity; white area: asynchronous irregular activity. (b) Top: firing rates from simulations (dots) and mean-field analysis (lines) for the excitatory (blue) and inhibitory (red) populations. Bottom: frequency of oscillations predicted from linear stability analysis (black line) and from simulations (dots) for *g* = 3.6. The gray area marks the parameter range where the network is in the synchronous irregular phase (i.e. where the eigenvalue is positive). (c) Examples of network dynamics from simulations. Left column: LFP amplitude; middle column: LFP autocorrelation function; right column: raster plot of 800 excitatory (top) and 200 inhibitory (bottom) neurons selected at random, with population firing rates of E (blue) and I (red) neurons. Rows correspond to *g* = 3.6 and *ν*_*ext*_*/ν*_θ_ = 1.3, 0.95, 0.65, respectively.

This behavior mirrors classic E–I networks with no synaptic dynamics [42], which exhibit a slow synchronous-irregular (SI) regime and an asynchronous irregular (AI) regime separated by Hopf boundaries as external drive increase. Slow oscillations are set by membrane and synaptic time constants and rely on recurrently shared fluctuations to align phases through the delayed E-I loop. As the external drive increases, a larger fraction of the variability of each neuron comes from the external (unshared) component rather than from recurrent interactions shared across the network. This shift effectively weakens the synchronization capacity of the circuit, lowering the effective loop gain of the recurrent feedback—and pushes the system away from the conditions that sustain the slow coherent mode, yielding an asynchronous-irregular state [42]. In contrast to instantaneous synapse models, here we include finite synaptic rise and decay in the recurrent dynamics; this synaptic filtering acts as a low-pass filter, shifting the resonance to lower frequencies and producing sustained, more sinusoidal slow oscillations. Consequently, the slow SI we observe is not organized into burst–quiescence cycles as in [42], but appears as a continuous 10–30 Hz activity, in line with motor-cortex data.

### Analysis of the model: spatio–temporal patterns

We now turn to the analysis of the network with spatially structured connectivity. To obtain the system’s dispersion relation, we solved numerically the characteristic equation (2) for |**k**| *>* 0. For isotropic connectivity, Fig. 4.a shows Re *λ*(*k*) as a function of the wavenumber magnitude *k* = |**k**| and of the external input strength *ν*_ext_, at the excitation–inhibition ratio used in Figs. 1–2 (see Table 1). There exists an interval of *ν*_ext_ for which Re *λ*(*k*) is positive at a finite wavenumber *k* > 0, indicating a spatio–temporal (Turing–Hopf) instability. For larger *ν*_ext_, Re *λ*(*k*) < 0 for all *k*, and the asynchronous, spatially homogeneous state is linearly stable. Fig. 4.b shows the most unstable wavenumber *k*^∗^ and its oscillation frequency as a function of *ν*_ext_, together with values computed from simulations; despite the finite network size, theory and simulations agree well. As expected, discrepancies increase at small *k* (long wavelengths) where finite-size effects are more pronounced (see Sec. 4.8). The role of delays is summarized in Fig. 4c, which displays Re *λ* as a function of *k* for different axonal conduction speeds *v* (equivalently, different distance-dependent delays). For large *v* (short delays), the dominant mode occurs at *k* = 0, consistent with a purely temporal Hopf instability and no spatial pattern. As delays increase (smaller *v*), the maximum shifts to *k*^∗^ *>* 0, revealing the emergence of spatio–temporal instabilities. The inset reports the monotonic dependence of *k*^∗^ on *v*. Figure 4d illustrates a representative plane-wave episode in the LFP amplitude from simulations within the unstable region; the raster below (random neuron subset) shows irregular spiking despite the mesoscopic wave. Figure 4e contrasts this with activity in the asynchronous regime. Because of sparse connectivity and finite-size fluctuations, spatio–temporal patterns still arise, but their morphology differs from planar waves: they form irregular structures of varying shape and spatial extent, often appearing patchy rather than coherent (see also Fig. S3).

**Figure 4:**
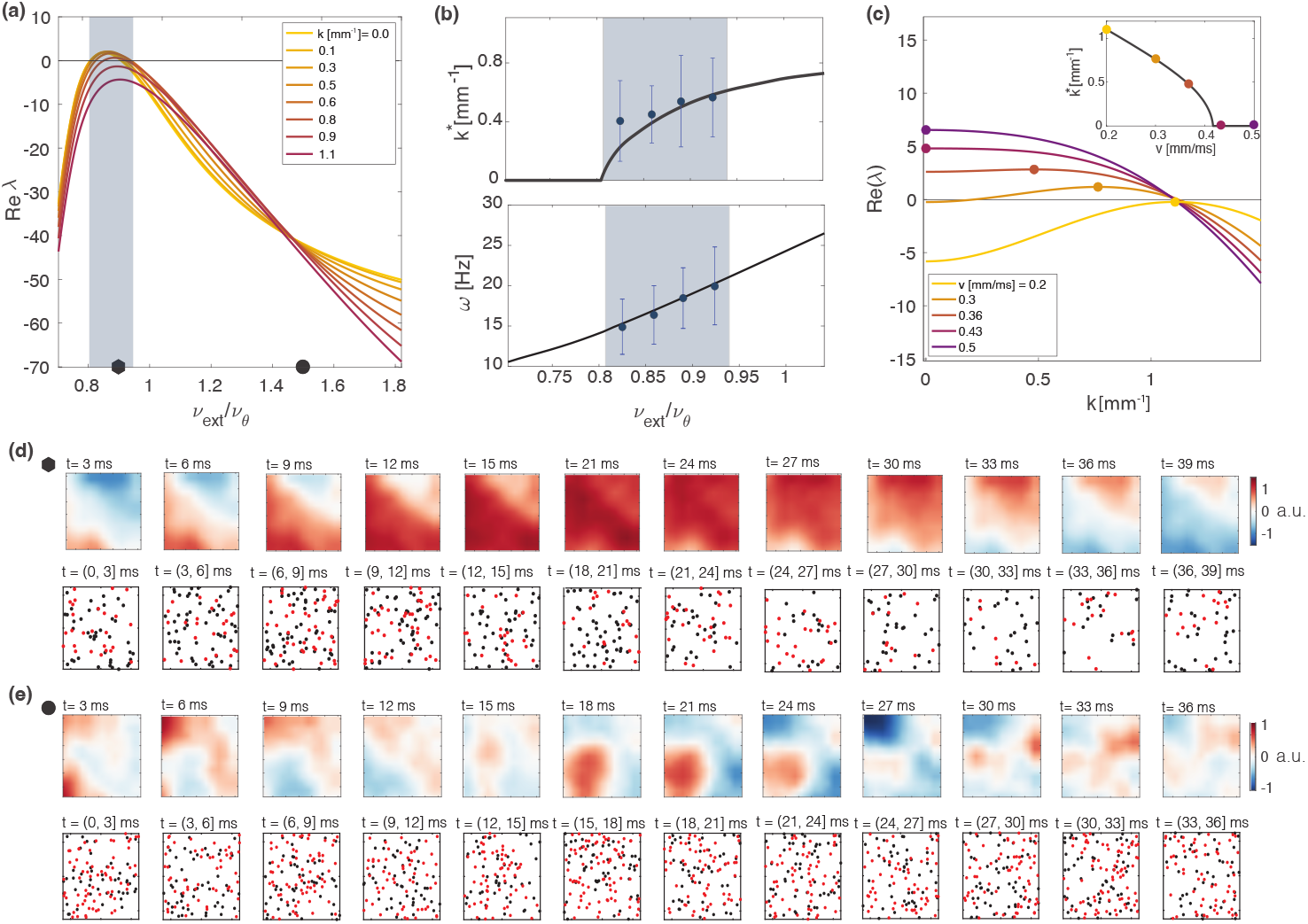
Spatio-temporal patterns of activity in the spatially extended model with isotropic connectivity. (a) Real part of the largest eigenvalue *Re*(*λ*) from the linear stability analysis of the spatially extended model, as a function of external input strength, for *g* = 4.1. Each colored curve corresponds to a different spatial wavenumber *k*. The gray area marks the region of spatio-temporal instability. (b) Top: wavenumber of maximal growth *k*^∗^, defined as the value of *k* that maximizes the real part of the leading eigenvalue, as a function of external input strength. Dots represent the mean wavenumber of wave events from simulations, with error bars indicating the standard deviation. Bottom: predicted frequency of oscillation of the spatio-temporal instability compared to the dominant frequency of oscillations of population rates from simulations (mean ± s.d.). (c) Real part of the largest eigenvalue *Re*(*λ*) as a function of the wavenumber *k* (maxima shown as dots). Different colors represent different propagation velocities of axonal delays in the model. Inset: wavenumber of maximal growth *k*^∗^ as a function of the propagation velocity. Dots in the main plot correspond to those in the inset. Here, *g* = 4.1 and *ν*_ext_*/ν*_θ_ = 0.35. (d) Example of a wave event detected from simulations, for parameters such that the system lies in the spatio-temporal instability regime (compare the symbol to panel c). Top: proxy of the LFP; bottom: spiking activity of a random subset of neurons (excitatory in blue, inhibitory in red). (e) Example of activity when the parameters are such that the homogeneous state is stable, where spatial gradients emerge due to heterogeneity in the connectivity structure and and finite-size fluctuations.

Importantly, qualitatively similar dispersion curves are obtained when distance-dependent delays are replaced by homogeneous delays while retaining spatially structured connectivity (Fig. S4). For sufficiently large delays, the maximum eigenvalue peaks at positive *k*, indicating that spatio–temporal instabilities can be present in the absence of spatially dependent delays. This shows that the instability arises intrinsically from the interplay between spatially structured coupling, synaptic filtering, and finite delays [20], rather than from passive phase propagation imposed by spatial delay gradients.

### Direction of wave propagation

Traveling waves detected from recordings propagate predominantly along a specific direction (Fig. 5e), corresponding to the rostro–caudal axis of the macaque motor cortex [3]. Classical anatomical studies have demonstrated that intrinsic horizontal connections in monkey area 4 are anisotropic, with the spread of degenerating fibers forming an ellipse whose long axis is anteroposterior (rostral–caudal) [44]. However, because axonal degeneration methods cannot identify the cells of origin or postsynaptic targets of individual fibers, it remains unclear whether these projections arise from pyramidal neurons or interneurons, and whether they preferentially innervate excitatory or inhibitory cells. More recently, Hao et al. [45] used microstimulation and multi-electrode recordings to show that evoked activity in the monkey motor cortex spreads anisotropically. This spread favors the rostro–caudal axis near the central sulcus and the medio–lateral axis at anterior sites, aligning with Rubino et al. [3]. However, these effective connectivity measures do not uniquely identify the underlying structural connectivity or specific cell types involved. Motivated by these findings, here we study how anisotropy in different connection types affects the direction of wave propagation in our model. Fig. 5 summarizes the model results: with isotropic connectivity (*ρ* = 0), waves can propagate in any direction, whereas introducing anisotropy breaks rotational symmetry and selects a dominant propagation axis. Panels b–c show the real part of the dominant eigenvalue, Re *λ*_max_, over the two-dimensional wavenumber plane, after rotating coordinates so that *u* and *v* denote the directions of maximal and minimal spatial spread of the connectivity, respectively. For isotropic connectivity (panel b), the contour of maximal Re, *λ*_max_ in (*u, v*)-space is circular, indicating no preferred orientation of the most unstable modes; consistently, simulations of the isotropic network (panel e) exhibit waves with approximately uniform propagation-angle distribution, up to finite-size fluctuations. When anisotropy is introduced in the E→E connections (panel c, *ρ* = 0.6), Re, *λ*_max_(*u, v*) develops two lobes centered at (*u, v*) ≈ (±*u*_0_, 0), elongated along the *v*-axis. These lobes identify symmetric clusters of unstable modes whose wavevectors have a dominant component along the *u*-direction, i.e. along the axis of maximal E→E spread; within each lobe a broad family of wavevectors with *u* ≈ ±*u*_0_ and small *v* are nearly equally unstable. As a result, simulations of the E→E-only anisotropic network exhibit waves that can propagate in many directions, but predominantly along the axis of maximal E→E spread (panel h).

**Figure 5:**
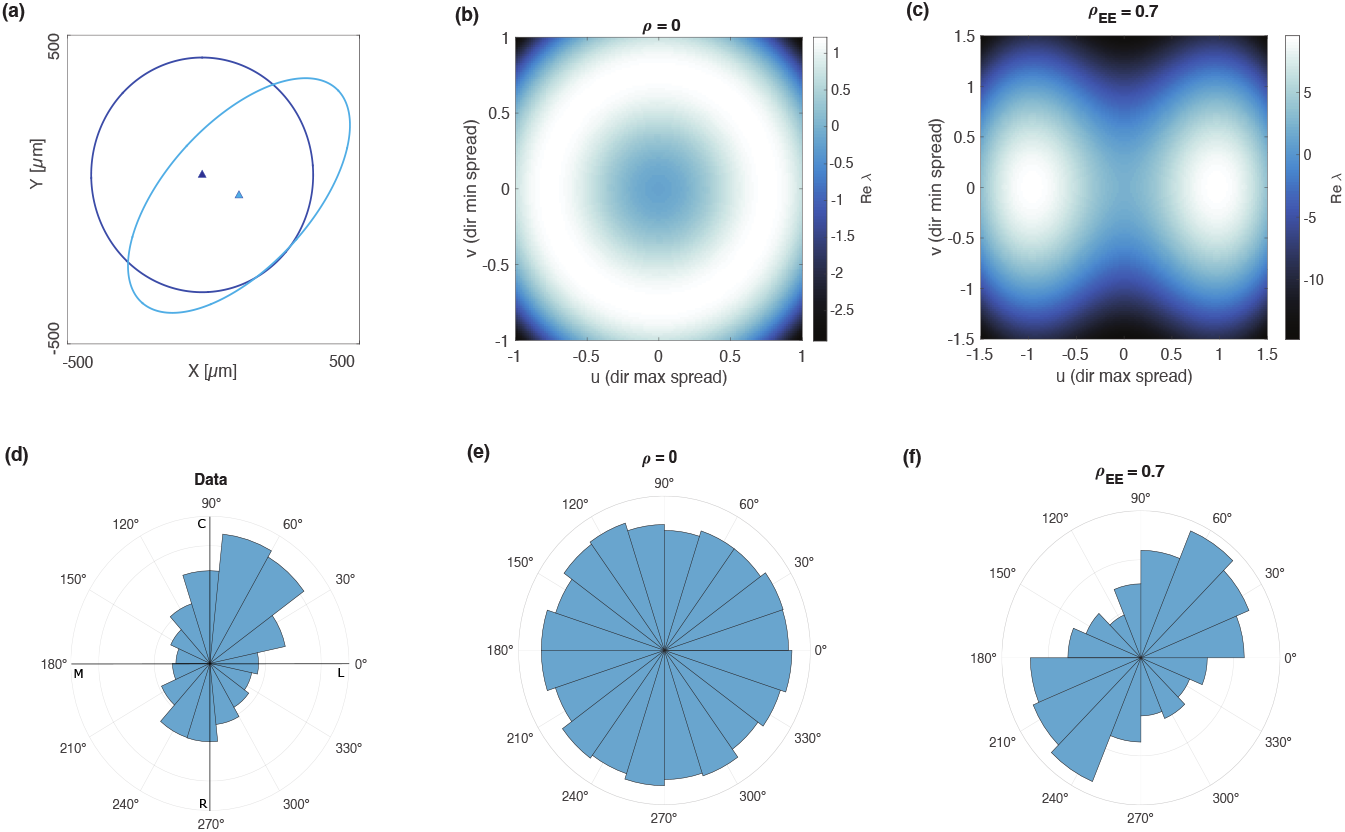
Effect of connectivity anisotropy on wave propagation direction. (a) Spatial spread of the E→E connectivity for the isotropic case (blue) and an anisotropic case (light blue). (b) Two-dimensional map of the dominant eigenvalue *λ*_max_(*u, v*) in the rotated wavenumber coordinates (*u, v*), where *v* corresponds to the axis of maximal connectivity spread and *u* to the axis of minimal spread, for a network with isotropic connectivity. (c) Same as (b), for a network with anisotropy in the E→E connections (*ρ* = 0.7). (e) Circular histogram of propagation directions derived from recordings. (f) Propagation directions from simulations with isotropic connectivity. (g) Same as (f), for a network with anisotropy in the E→E connections (*ρ* = 0.7).

In summary, the model shows that anisotropic E→E connectivity is sufficient to reproduce the rostro–caudal propagation bias observed experimentally. This provides a parsimonious circuit-level explanation for the directional structure of motor-cortical traveling waves, consistent with classical anatomical evidence for anisotropic horizontal connectivity. Figure S5 further shows that waves follow the direction of maximal spread for E→E or I→I anisotropy alone, but shift to the direction of minimal spread when anisotropy involves both excitatory and inhibitory pathways.

We note that Figs. 1 and 2 were performed using a model in which the E→E connections are anisotropic (*ρ* = 0.6, see parameters in Table 1).

## 3 Discussion

In this work, we show that a spatially structured LIF network with realistic synaptic kinetics recapitulates the spatiotemporal organization of beta-band activity observed in the macaque motor cortex during instructed reaching [3]. Specifically, our model reproduces the transition from traveling waves during motor preparation to spatially graded beta attenuation at movement onset, together with the irregular activity characteristic of movement execution. The model further predicts that anisotropic long-range excitatory-to-excitatory connectivity can account for the rostro–caudal directional bias of propagating beta waves. Mechanistically, beta oscillations emerge from delayed negative feedback within the local E–I loop, driven by the interplay of membrane time constants, conduction delays, and synaptic kinetics, with inhibition operating on a slower timescale than excitation. These oscillations manifest as traveling waves via a Turing–Hopf instability of the spatially homogeneous state. Notably, our model demonstrates that while spatially dependent conduction delays are not strictly required for this instability to emerge, they must exceed a threshold to support coherent wave propagation. The network shifts from an oscillatory to an asynchronous regime as external excitatory drive increases, modeling external inputs arriving in the motor cortex prior to movement onset. This interpretation is supported by recent causal evidence showing that time-varying thalamocortical and midbrain–thalamocortical inputs are required to initiate movement in mice [46]. In the asynchronous regime, randomness in the connectivity, finite-size fluctuations, and transmission delays still give rise to irregular, non-planar spatiotemporal patterns, reminiscent of the traveling waves reported in balanced cortical network models operating in the asynchronous state [26].

A recent rate-based model of motor cortex has shown that local E–I modules coupled by long-range excitation can reproduce transient beta oscillations and traveling waves during movement preparation [27]. In that framework, local E–I loops are resonant in the beta range but the circuit operates near but below the oscillatory instability; stochastic inputs dephase otherwise synchronized modules, and waves reflect this dephasing process. Our spiking model builds on related principles but implements a fully spatially extended E–I network with sparse, distance-dependent recurrent connectivity and conduction delays in all pathways. In this framework, we identify a Turing–Hopf instability at finite wavenumber, so that in the mean-field limit the dominant eigenmode is a self-sustained traveling beta wave; when the network operates close to this bifurcation, sparse connectivity and finite-size fluctuations disrupt the coherent unstable mode, producing intermittent wave bursts. Taken together, the two models can be viewed as subthreshold and superthreshold realizations of the same underlying E–I wave-generating mechanism. The two models also make related predictions about directionality: when anisotropic long-range excitation targets both E and I populations, waves propagate along the axis of minimal effective spread, consistent with [27], who therefore attributed the rostro–caudal propagation bias in M1 to anisotropic inputs. Our pathway-specific analysis shows, however, that if anisotropy is restricted to the E → E pathway, the most unstable modes instead align with the axis of maximal spread, providing a parsimonious circuit-level explanation for the rostro–caudal directionality bias observed in motor cortex. Whether this bias indeed reflects anisotropic local connectivity, or instead arises from anisotropic external inputs, is a testable prediction that future experiments targeting cell-type–specific horizontal projections and the spatial distribution of thalamocortical or premotor inputs to M1 could resolve.

Beyond the task-related modulations explored here, our phase diagram shows that changes in local E/I balance can reposition the circuit relative to the beta-generating instability. In particular, decreasing the relative strength of inhibition (smaller *g*) moves the operating point deeper into the oscillatory regime, making the beta mode more robust (Fig. S2b). Speculatively, this suggests that perturbations that weaken cortical inhibition could promote abnormally persistent beta oscillations. Exaggerated and overly synchronized beta-band activity in cortico–basal ganglia circuits is a well-established hallmark of Parkinson’s disease and is causally linked to bradykinesia in both patients and animal models [47, 48, 49, 50]. Although our model does not include the basal ganglia–thalamo–cortical loop or dopamine-dependent plasticity, it provides a local-circuit mechanism by which shifts in E/I balance could render motor cortex more susceptible to sustained beta entrainment, potentially contributing to the amplification of pathological beta rhythms in parkinsonian states.

While we focused here on the emergence and attenuation of beta oscillations, preliminary simulations revealed additional phenomena that we did not explore in detail (see Supplementary Fig. S6). First, when the external drive is close to the lower boundary of the oscillatory regime, where mean firing rates are very low, the network exhibits a low-beta rhythm consistent with our linear stability analysis, but the LFP power spectrum also displays a secondary peak in the alpha range. This additional timescale may reflect nonlinear interactions and/or finite-size fluctuations that are not captured by the dominant beta mode in the linear theory. Clarifying the origin of this alpha activity, and its interaction with beta, would be particularly interesting given experimental reports of coordinated alpha–beta dynamics in sensorimotor circuits [51, 52]. Second, for external inputs that drive the network well above the asynchronous regime, the model develops gamma-band oscillations dominated by the inhibitory feedback loop. This behavior parallels the fast oscillatory instability described in random LIF networks by Brunel [42], where strong recurrent inhibition combined with finite transmission delays gives rise to high-frequency rhythms. Future work could build on our modeling framework to investigate how spatially structured gamma-band patterns emerge from circuit properties. This is particularly relevant in light of recent experimental work showing that single-trial propagation directions of high-gamma recruitment in M1 are movement-specific and carry kinematic information, whereas beta-band traveling waves express stereotyped rostro–caudal propagation and convey condition-independent signals related to movement initiation rather than movement details [53]. Understanding why beta waves exhibit stable propagation patterns while gamma recruitment directions flexibly rotate with movement parameters could provide a deeper view of how different frequency bands support distinct functional roles in motor cortical computation.

The computational role of cortical traveling waves is an active topic of research. Proposed functions include dynamically gating or routing information between cortical areas via coherence-based communication [54, 55], imposing mesoscopic temporal reference frames for organizing local population codes [56], providing a spatiotemporal context that modulates local excitability and coordinates spiking across micro-to macroscopic scales [2, 1, 57], implementing sequential sampling or predictive sweeps over internal representations within predictive-processing frameworks [58], and supporting nontrivial visual computations such as image segmentation in recurrent network models [59]. In behaving primates, spontaneous low-frequency waves gate sensory detection, with perceptual sensitivity and neuronal responses depending on wave phase at stimulus onset [25], and human intracranial recordings indicate that wave direction and phase systematically switch with cognitive regime, consistent with distinct wave modes indexing different computational states [60]. Intracranial recordings in patients recovering from traumatic brain injury further show that the re-emergence of beta-band traveling waves over cortex closely tracks neurological recovery and predicts survival, suggesting that such waves index the restoration of large-scale integrative dynamics [61]. In motor cortex, recent work has provided causal evidence that propagating beta-attenuation patterns are necessary for movement initiation: patterned intracortical microstimulation delivered against the natural propagation direction selectively delays reaction times, while stimulation with the natural direction does not [4]. Moreover, perturbing the spatial order of beta amplitudes along the propagation axis degrades muscle electromyographic decoding more than orthogonal perturbations, suggesting downstream circuits are sensitive to the cortical recruitment order imposed by these waves. Our model suggests that the spatial order of motor recruitment may arise as a direct consequence of the motor cortex’s intrinsic E–I architecture and connectivity anisotropy, providing a mechanistic framework for interrogating the causal and computational roles of propagating beta waves in motor control.

## 4 Methods

### 4.1 Model

Let *a* ∈ {*E, I*} label the population, and let *i* index a neuron in that population with position 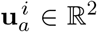 on the cortical sheet. The time course of the membrane potential 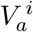 of each neuron *i* belonging to population *a* ∈ {*E, I*} is modeled as

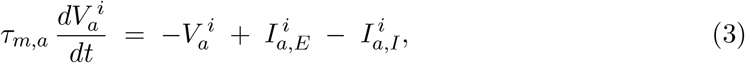

where the currents 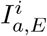 and 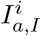 represent, respectively, AMPA-receptor mediated and GABA-receptor mediated currents. The time scale of integration of input currents is defined by the membrane time constant *τ_m,a_*. When the voltage reaches the spike threshold, a spike is emitted and the voltage is kept fixed at a reset value during a short refractory time 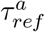. AMPA synaptic currents are triggered by spikes from excitatory pre-synaptic neurons (both from outside and inside the network), while GABA synaptic currents are generated by spikes from inhibitory pre-synaptic neurons. The time course of post-synaptic currents is described by a delayed difference of exponentials, with different time constants for AMPA/GABA mediated pre-synaptic currents, similar to previous network studies [28, 62, 63]. Such time courses are obtained through auxiliary synaptic variables 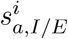 that obey the following system of equations:

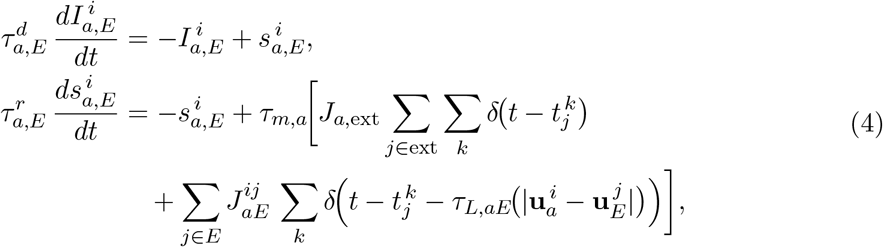

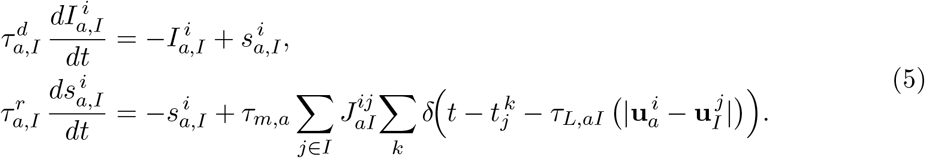

where 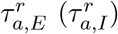 is the rise time of the synaptic AMPA (GABA) currents, while 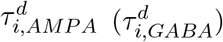 is the decay time of the synaptic AMPA (GABA) currents. External spikes are generated through an inhomogeneous Poisson process with rate

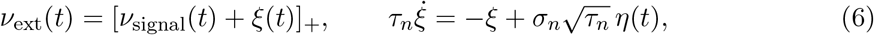

with *η* Gaussian white noise, where the filtered noise term *ξ*(*t*) is motivated by the need to capture the low-frequency components of the LFP power spectrum observed in data (see [62, 63]). The parameter *τ_n_* denotes the correlation time of the noise process *ξ*(*t*), which sets the timescale over which the external input fluctuates. Synaptic delays 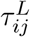 depend on the distance between the location of pre- and postynaptic cells, respectively 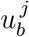 and 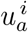:

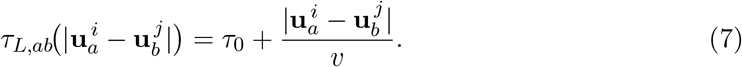

where *τ*_0_ is a distance-independent delay, and *v_c_* is the axonal conduction speed; we assume that axonal conduction speed is isotropic, so *τ_L,ab_* depends only on the Euclidean distance between cells.

Synaptic efficacies from an external, excitatory, or inhibitory presynaptic neuron *j* ∈ {ext, *E, I*} onto a postsynaptic neuron *i* ∈ {*E, I*} are denoted by *J*_a,ext_, 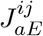, and 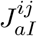, respectively. Synaptic connectivity is random with a distance dependent probability. We set 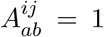 if neuron *j* ∈ *b* connects to neuron *i* ∈ *a* (and 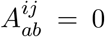 otherwise), with connection probability

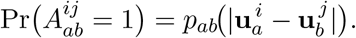

The synaptic efficacy is

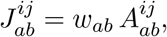

where synaptic weights *w_ab_* take a fixed value determined by the presynaptic population *b* and postsynaptic population *a*. The analysis that follows applies to any connection probability *p_ab_* that is integrable, nonnegative, normalized as above, and has a well-defined Fourier transform. In this work, we use an elliptical exponential kernel with varying degrees of anisotropy. If 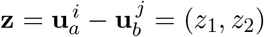 denotes the vector connecting the location of a neuron in population *a* and one in population *b*, then:

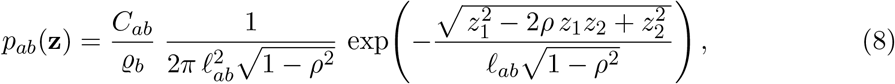

*ℓ*_ab_ *>* 0 being the spatial spread of the connectivity, and *ρ* representing the degree of anisotropy, |*ρ*| *<* 1. The above connection–probability profile is normalized so that so that

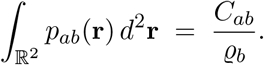

where *C_ab_* is the average number of connections from population *b* to *a* and *ϱ_b_* the areal density of population *b*.

All model parameters are listed in Table 1.

### 4.2 Recording grid and LFP proxies

We simulated a square cortical sheet of side *L* = 2 mm. The sheet contained a surface density of 2.8 × 10^4^ neurons*/*mm^2^, corresponding to a total of *N* ≃ 1.12 × 10^5^ neurons partitioned into *N*_E_ = 0.8 *N* excitatory cells and *N*_I_ = 0.2 *N* inhibitory interneurons. To mimic an electrode array, we placed *N*_R_ recording sites on a regular square grid with spacing *d*_grid_ = 400 *µ*m, leaving a 200 *µ*m margin from each edge to minimize boundary effects, for a total of *N*_R_ = 25 recording sites. For Figs. 4-5, we use a denser array with *d*_grid_ = 140 *µ*m, and also leaving a 200 *µ*m margin from each edge, for a total of 144 recording sites.

To compute a proxy of the local field potential (LFP) from our point–neuron network, we adopted the approach of [40, 64, 39], which showed that the LFP generated by a three-dimensional network model of multi-compartmental neurons with realistic morphology can be approximated by a linear combination of excitatory and inhibitory synaptic currents onto excitatory neurons (for a review, see [65]). In this formulation, the single–neuron contribution to the LFP factorizes into a purely time–dependent and a purely distance–dependent component —a proxy that accounts for over 90% of the variance of the detailed LFP signal across different depths and recording positions [39]. Following, [39], the temporal component of the LFP at electrode *i* is modeled as the sum of the amplitude of the AMPA and GABA synaptic currents from nearby pyramidal neurons:

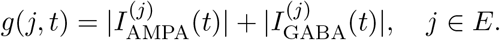

For the spatial component, we model the distance-dependent attenuation with a shape function, *f* (*r*), where *r* represents the radial distance from a contributing neuron. This function, introduced by [40], captures both the near-field and far-field decay of the single-cell contributions to the LFP. The spatial separation between apical and basal current sinks and sources in pyramidal neurons produces a dipole-like field with a far-field 1*/r*^2^ decay. At small neuron–electrode separations, the decay is shallower because the extracellular potential reflects spatially distributed transmembrane currents along the dendrites; the dendritic cable acts as a frequency-dependent low-pass filter (in time) and a spatial smoother, so high-frequency components are more strongly attenuated and the fall-off is slower than the ideal dipolar law. Accordingly, we use the piecewise form in Eq. (9), which captures both regimes, where *r_x_* denotes the crossover distance between them:

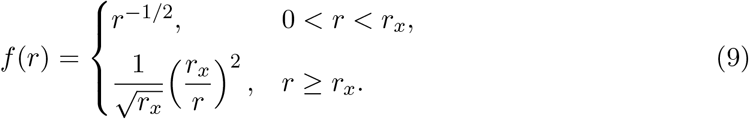

Biophysical modeling indicates that *r_x_* for layer 2/3 pyramidal neurons is typically around 100–150 *µ*m when neurons receive uncorrelated inputs, and decreases with frequency [64, 40]. We fixed *r_x_* = 130 *µ*m as a representative low–frequency value. For computational tractability, we limited the spatial summation to neurons within the 95% spatial reach of the model, corresponding to *R_r_* ≈ 1.85 *r_x_* ≃ 278 *µ*m, which retains nearly all of the theoretical LFP contribution while reducing runtime. The resulting (unnormalized) LFP at electrode *i* is

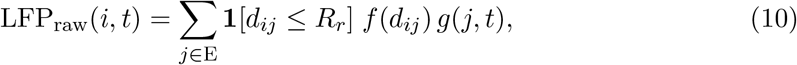

where *d_i,j_* denotes the Euclidean distance between recording site *i* and neuron *j*, and **1**[·] is the indicator function limiting the spatial summation. We normalize by the total spatial weight to remove dependence on the number of contributing cells:

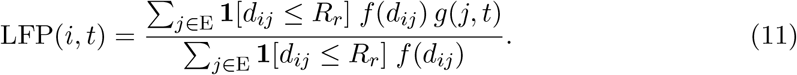

### 4.3 Diffusion approximation

We turn to a continuous spatial description with coordinates *x* ∈ ℝ^2^. The summed synaptic shot noise is approximated by colored Gaussian noise in the limit of small synaptic couplings *w_ab_* [42]. In this approximation, the dynamics of a neuron *i* in population *a* is given by:

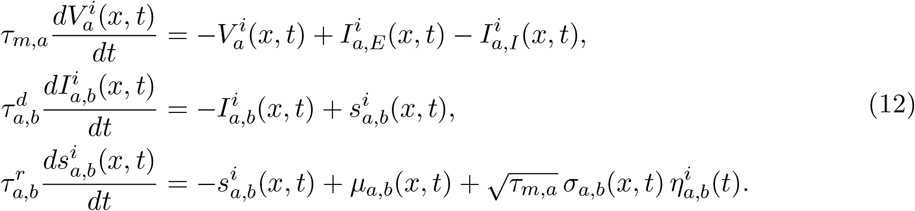

with *η_a,b_* independent white-noise processes; *µ_a,b_* and 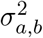 denote the mean and variance of the shot-noise drive entering *s_a,b_*. Because the axonal delay depends on distance, the presynaptic rate enters the mean input drive via a delayed spatial integral:

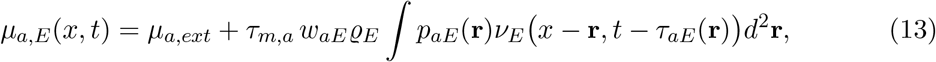

and similarly for the input variances,

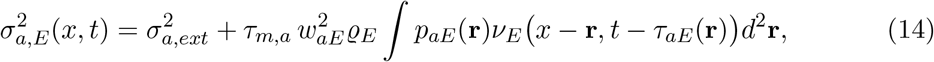

where the external input contributions are

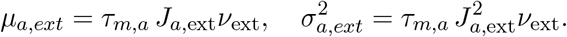

Analogous expressions to (13-14) hold for GABAergic input, except that the external drive contribution is absent.

#### Homogeneous stationary state

For our analysis, let us focus on the case of constant rate of external input: *ν*_ext_(*t*) = *ν*_ext_. The spatially homogeneous, stationary rates 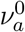 are obtained via the effective-noise reduction of Fourcaud and Brunel [66], which maps the high-dimensional Fokker-Planck system – accounting for the joint distribution of {*V_a_, I_a_, s_a_*} – under colored synaptic noise to an equivalent LIF with a threshold shift proportional to 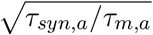. Defining the effective total input

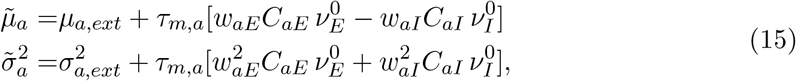

the stationary rate

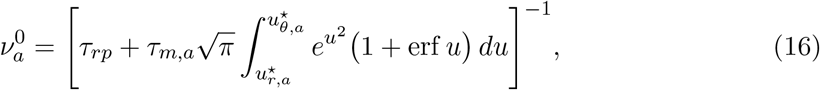

with the colored–noise–corrected bounds

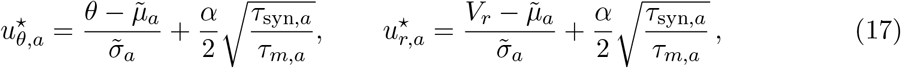

is valid to first order in 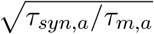. The constant *α* is 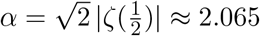, *ζ* denoting the Riemann zeta function. The effective synaptic time constants *τ_syn,a_* are given by:

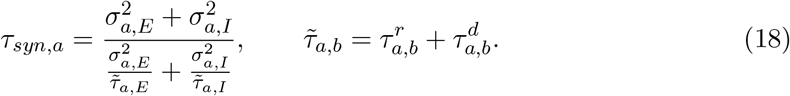

The coefficient of variation (CV) of the interspike interval can be also computed following [42] and using the Fourcaud–Brunel effective-threshold shift:

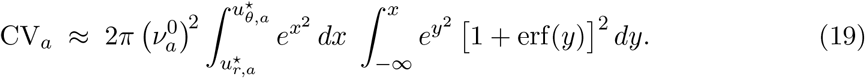

#### Threshold external rate under colored synaptic noise

External inputs in plots are reported normalized by *ν_θ_*, defined as the external rate required to reach threshold in the absence of recurrent input, taking into account the effective threshold shift due to colored synaptic noise. With *τ_syn,a_* given by Eq. (18):

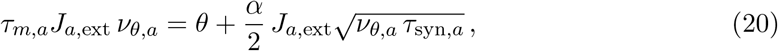

giving:

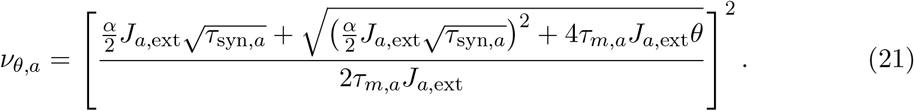

For the plots, we considered the population average *ν_θ_* = 0.8*ν_θ,E_* + 0.2*ν_θ,I_*.

### 4.4 Linear stability analysis

We introduce a small spatio-temporal perturbation to the homogeneous and stationary presynaptic firing rate:

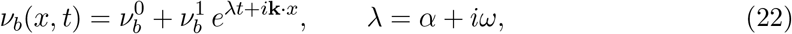

with 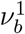 small. Inserting into the equations (13)–(14) for the mean and variance of the raw synaptic drive, we get:

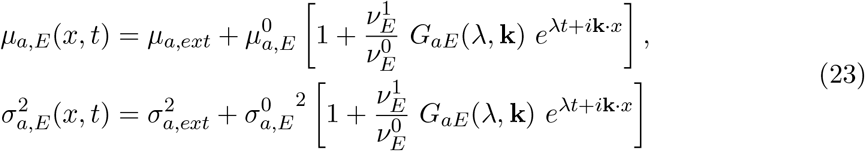

where

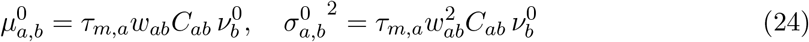

are the mean and variance of the *recurrent* homogeneous and stationary input drive from population *b* to *a*. Eq (23) represents AMPA input drives (*µ_a,E_*, 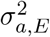); the mean and variance of the GABA input drive (*µ_a,I_*, 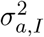) has analogous form, except that the contribution of external inputs is absent. The spatio-temporal kernel is:

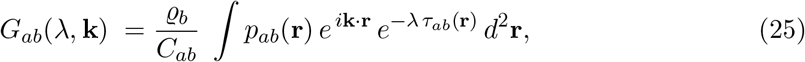

where the spatially-dependent delay was defined in (7). For constant delays, *τ_ab_*(**r**) ≡ *τ_L_*, this reduces to

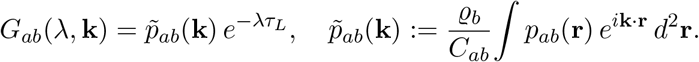

Applying the synaptic filters to (23) yields the following modulations of the postsynaptic current mean and variance:

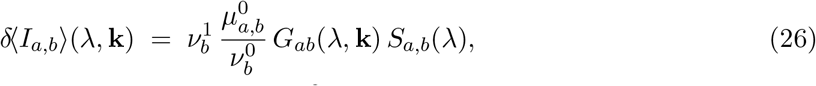

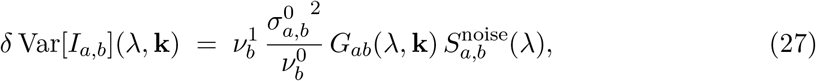

with synaptic filters:

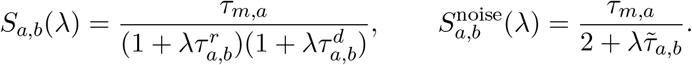

Note that, for the variance, we approximate the rise and decay filters by a single effective time constant (18), following [66].

To relate these modulations to the postsynaptic firing rate, we linearize around the operating point and write the first-order response:

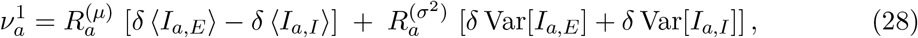

where 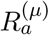 and 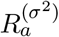 are the susceptibilities to perturbation in the mean and variance, respectively. By requiring the postsynaptic perturbation (28) to match the presynaptic one (22), we obtain the 2D eigenvalue problem:

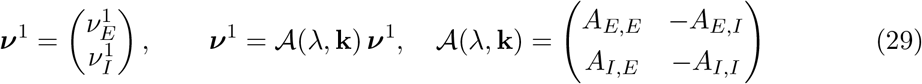

where we have defined:

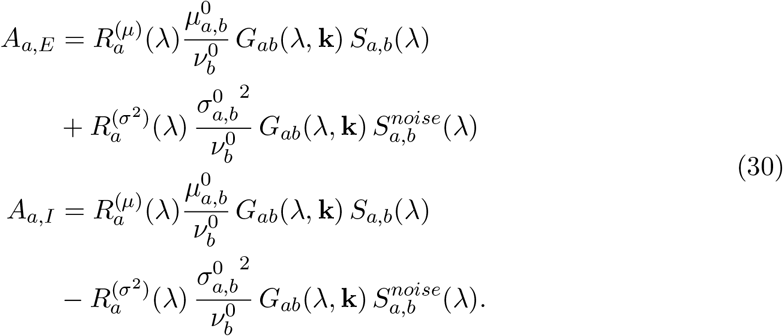

Nontrivial solutions to (29) exist if the characteristic equation (2) is satisfied.

### 4.5 Spatio-temporal kernel and direction of wave propagation

The spatio-temporal coupling kernel *G_ab_*(*λ*, **k**) defined in (25) is derived from the elliptically symmetric distance-dependent connectivity profile (8) combined with distance-dependent axonal delays (7), allowing for anisotropic connectivity through a correlation parameter *ρ*. In the isotropic case, (*ρ* = 0), *G_ab_*(*λ*, **k**) is evaluated analytically; for the anisotropic case (*ρ >* 0), we compute the kernel numerically to determine the dispersion relation and the direction of maximal instability. Full derivations and numerical procedures are provided in SI Appendix.

### 4.6 Single-Neuron Response Functions

The linear response of individual neurons to perturbations in input mean and variance is defined by the susceptibility functions 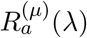 and 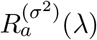. Analytical expressions exist for white-noise–driven leaky integrate-and-fire neurons [41, 67]. To incorporate finite synaptic filtering in 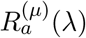, we construct colored-noise corrections by combining analytical asymptotic results for mean perturbations [66] with numerical simulations of single LIF neurons, yielding a fitted mapping that transforms white-noise susceptibilities into their colored-noise counterparts across the parameter space. The same strategy is extended to perturbations of the input variance under colored noise, a regime which, to our knowledge, has not been treated analytically. Detailed derivations, simulation protocols, and fitted parameter values are described in the SI Appendix, together with the resulting plots for 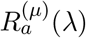 and 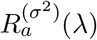 (Fig. S1).

### 4.7 Wave detection and kinematics from recordings and simulations

We identified the dominant frequency *f_dom_* from a Welch power spectral density of the spatial-mean LFP, and extracted the phase in the narrow band [*f_dom_* − Δ*f, f_dom_* + Δ*f*], with Δ*f* = 3 Hz. All channels were band-pass filtered in this band (4^th^-order Butterworth) and converted to analytic signals *V* + *i* ℋ{*V*} = *a e^iϕ^* with the Hilbert transform ℋ{.}, yielding instantaneous phase maps *ϕ*(*x, y, t*) on the electrode grid with spacing *d_x_*; phases were spatially unwrapped in 2D on each frame (reference-centered) prior to computing spatial gradients.

To identify traveling waves, we required that two complementary criteria were met simultaneously. First, we quantified how consistently local phase gradients were aligned across the array using the Rayleigh statistic *Z*. For each time point, we computed the spatial phase gradient ∇*ϕ*(*x, y, t*) and normalized it to a unit vector **u**(*x, y, t*) = ∇*ϕ/*∥∇*ϕ*∥. The mean of these unit vectors, **ū**(*t*) = ⟨**u**(·, ·, *t*)⟩*_x,y_*, represents the dominant orientation of phase flow at that time. We then defined

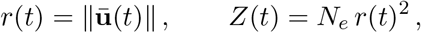

where *N_e_* is the number of electrodes, and the norm ∥ · ∥ denotes the mean resultant length. *Z*(*t*) quantifies how strongly gradient directions align across the array: large values indicate that phase gradients are well aligned across space, consistent with a plane wave. Second, we assessed how well the instantaneous phase map *ϕ*(*x, y, t*) was fit by a plane wave 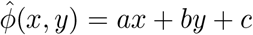 using least-squares regression. For each time point we computed the root-mean-square residual,

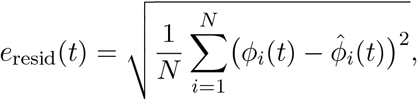

where *N* is the number of electrodes. To obtain a null distribution for statistical thresholds, we generated conservative spatial surrogates at each time by randomly shuffling electrode locations in the phase map, recomputed the Rayleigh statistic *Z* and the plane-fit residual on these surrogates, and pooled values over time and trials to form empirical nulls. Wave epochs were defined as contiguous time intervals of at least 10 ms in length where both *Z*(*t*) exceeded the 99.99-th percentile of the pooled surrogate *Z* distribution and the plane-fit residual fell below the 20-th percentile of its surrogate distribution. We adopted the Rayleigh-*Z* alignment metric with surrogates because its percentile-based thresholds are stable across different array sizes (here 10 × 10 in data vs 5 × 5 in simulations). The plane-fit condition was included to allow direct comparison of the measured wave kinematics with our theoretical predictions from linear stability analysis, where instabilities manifest as perturbations of the form *e*^*λt+i***k**·**x**^, that is, spatiotemporal modes with an ideal plane-wave structure.

Wave kinematics followed phase conservation, ∂_*t*_*ϕ* + **v**·∇*ϕ* = 0, which gives the instantaneous speed estimate

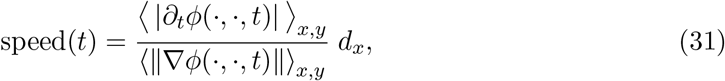

where ∂_*t*_*ϕ* is computed by finite differences (Δ*t* = 1 ms). For each detected wave we averaged ∇*ϕ* over space and time to obtain a single direction vector; the propagation direction was taken as the direction of decreasing phase, −⟨∇*ϕ*⟩*_x,y_*; to resolve the *π*-ambiguity, we oriented the angle using the sign of the mean temporal phase slope. Wavelength was computed from the mean gradient magnitude,

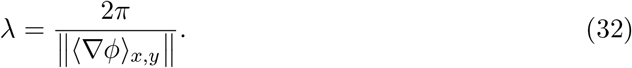

To ensure reliable quantification of wavelength, the electrode array should ideally span several wavelengths of the oscillation. In our simulations neurons were placed on a square sheet with edge *L* = 2 mm. Given that the expected wavelengths are on the order of 6 - 30 mm, the array size is below the detection threshold. This limitation explains the large error bars in Fig.3d, although the agreement with the theoretical prediction remains robust.

#### Beta attenuation times

For the analysis of *β*-attenuation times shown in Fig.2d–e (recordings) and Fig.2f–g (simulations), we use analogous procedures but defined the perievent window according to the experimental condition. In the recordings (Fig. 2d–e) the window spanned −400*/*200 *ms* relative to Movement onset, as in previous studies [4]. In the simulations (Fig. 2f–g) the window spanned −100/500 *ms* relative to the onset of the sigmoidal increase of the external input. Within each window we band-pass filtered the single-trial LFPs in the subject-specific *β*-band (*f*_dom_ ± 3 Hz) and extracted their analytic signals with the Hilbert transform to obtain amplitude envelopes; for the recordings the single-trial envelopes were first denoised using an auto-encoder. For each single trial the denoised *β*-band envelope was converted to normalized amplitude values, and the latency of *β*-attenuation was estimated using a threshold-crossing criterion.

### 4.8 Comparing mean field analysis and simulations of a finite system

Our linear stability analysis is derived for an infinite sheet, where the the linearized stability operator (the spatio-temporal convolution operator) is translation-invariant and thus diagonal in the spatial Fourier basis. Simulations, however, are performed on a finite square of side *L* = 2 mm with hard boundaries, which (i) discretizes the set of admissible wavenumbers and (ii) breaks exact diagonalization due to boundary-induced inhomogeneity. If the spatial kernel range *ℓ* and the delay length *v/ω* satisfy max{*ℓ, v/ω*} ≪ *L*, boundary effects are exponentially small in *L/ℓ*, and the bulk prediction remains accurate. In our case, *L* = 2 mm, *ℓ* = 0.38 mm, *v* = 0.3 mm*/*ms, and *ω* ≈ 20 Hz, so *v/ω* ≈ 2.4 mm is comparable to *L*. While our theory relies on several approximations, Fig. 4d demonstrates good agreement between the wavelength and oscillation frequency predicted by the theory and those observed in simulations, indicating that the approximation is sufficiently accurate.

For the spatially homogeneous model of Fig. 3, delays are computed as *τ_L,ab_* = *τ_0,ab_* + 𝔼*_ρ_*[∥**r**∥]/*v_ab_*, where the average 𝔼*_ρ_*[∥**r**∥*_ρ_*] is computed numerically by applying the connectivity mask on a finite 2 mm × 2 mm domain for better comparison with simulations, yielding the average delays 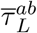 in Table 1.

### 4.9 Data

The data consist of electrophysiological recordings from the primary motor cortex of macaque Rk (141 units, 391 trials), as originally reported in [3]. For full details on the recording protocol and multi-electrode array implantation, see [3].

## Acknowledgments

We thank Bard Ermentrout, John Reynolds, Lyle Muller and Vincent Hakim for discussions on early versions of this model. This work was supported by grant R01NS104898.

## 5 Supplementary Information

### 5.1 Spatio-temporal kernel and direction of wave propagation

The elliptically symmetric exponential coupling density (8) can be written as

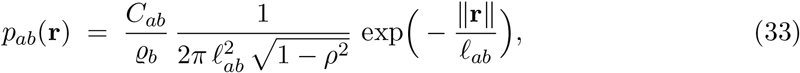

With 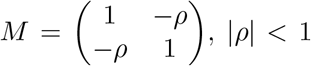, and 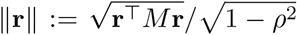, where |**r**| denotes the Euclidean distance between pre- and postsynaptic cells. The normalization holds on an infinite sheet. The distance-dependent delays are defined as

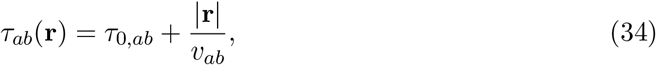

with direction-independent conduction speed *v_a,b_*. In the isotropic case (*ρ* = 0), the probability of connection (33) is radial and the spatio-temporal kernel (25) admits a closed-form expression:

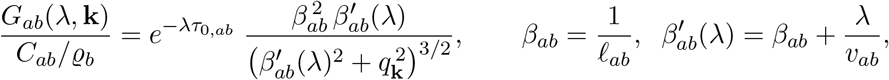

Where 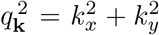. For anisotropic connectivity (*ρ >* 0), the combination of an elliptic *p_ab_* with Euclidean distance–dependent delays precludes a simple analytic form, and we evaluate the integral in Eq. (25) numerically.

In Fig. 3 we analyze a spatially homogeneous network; the kernel reduces to *G_ab_* = *C_ab_*/ϱ*_b_*, and the eigenvalue depends only on frequency, *λ* = *λ*(*ω*). In this case, we use the effective delay

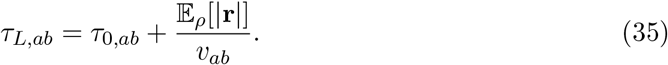

In Fig. 5 we consider anisotropic connectivity (*ρ >* 0) and analyze the dispersion relation in the rotated wavenumber coordinates

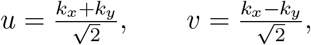

so that for *ρ >* 0 the *u*-axis corresponds to the direction of maximal spatial spread of the connectivity and the *v*-axis to the direction of minimal spread. We then plot the real part of the dominant eigenvalue, Re *λ*(*u, v*), over the (*u, v*) plane to characterize the locus and anisotropy of unstable modes.

### 5.2 Single-Neuron Response Functions

For a leaky integrate-and-fire (LIF) neuron driven by input with baseline statistics 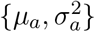 and no synaptic filtering, the linear responses to small sinusoidal perturbations of the input mean and variance were derived in [41, 67]:

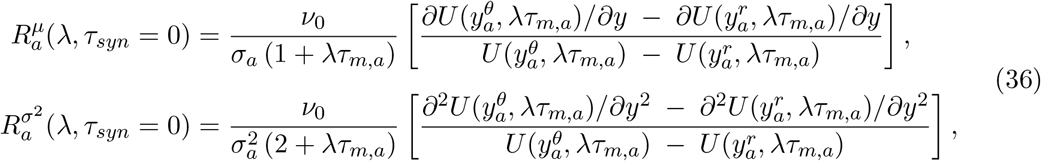

where 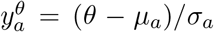 and 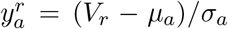 are the normalized threshold and reset potentials, respectively. The auxiliary function *U* is given by

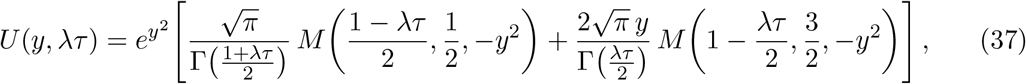

where *M* (·, ·, ·) is the confluent hypergeometric function and Γ is the Gamma function. In a model with synaptic filtering, the susceptibility to mean perturbations was analyzed in [68, 66], yielding exact small- and large-*ω* limits. At low frequencies (*ωτ_syn_* ≪ 1), the first-order corrections to the response amplitude due to synaptic filtering vanish [66]; consequently, the response reduces to the static susceptibility, given by the derivative of the stationary transfer function:

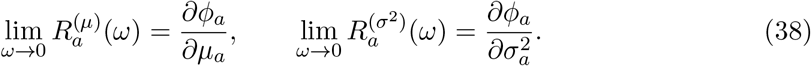

In contrast, at high frequencies, the synaptic time constant fundamentally changes the response scaling. The response amplitude approaches a plateau:

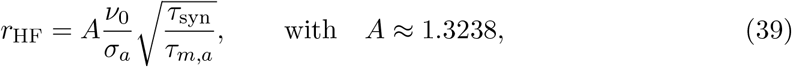

with a negligible phase lag. We obtain the full colored-noise response by simulating the LIF neuron and fitting a mapping that transforms the white-noise susceptibilities into their colored-noise counterparts across the parameter space. We extend this approach to perturbations of the input variance under colored noise, a regime which, to our knowledge, has not yet been treated analytically.

All colored-noise corrections are fitted on the imaginary axis, while retaining the exact *λ*-dependence of the white-noise susceptibilities. This approximation neglects explicit ℜ*λ*-dependence of the colored-noise correction and is accurate when |ℜ*λ*| *τ_syn_* ≪ 1. To numerically obtain the susceptibilities, we simulated a population of uncoupled neurons with colored input and apply oscillatory perturbations to either the mean or variance of the input current. For mean perturbations, the dynamics is governed by

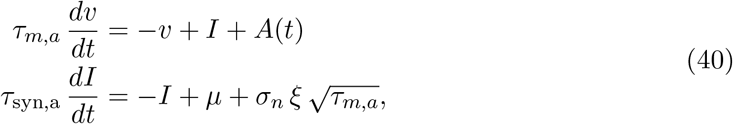

with stationary drive *A*(*t*) = *A*_0_ until equilibration, followed by oscillatory input *A*(*t*) = *ϵ* sin(2*πf*_ext_*t*) with *ϵ* = 0.05 *µ*. For variance perturbations, we use

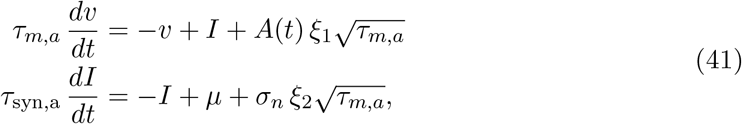

with *ϵ* = 0.2 *σ_n_* and independent standard white noise processes *ξ*_1_, *ξ*_2_. A refractory period *τ_rp_* was included, as in the full model. The amplitude and phase of the population rate are extracted via Fourier analysis. Simulations are performed for both excitatory and inhibitory neurons, using effective synaptic time constants *τ*_syn,a_ from Eq. (18). We repeat simulations at fixed network parameters for three values of the external Poisson rate *ν*_ext_ (see Fig.S1). Defining

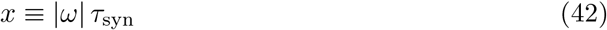

we fit the following functional forms simultaneously across the three *ν*_ext_ values, using the white-noise responses and asymptotic constraints:

#### Mean perturbation - amplitude

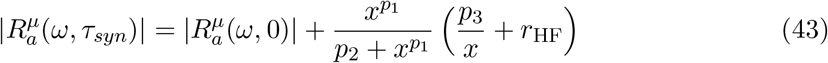

where 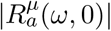 is the amplitude of the white noise response (36) when evaluated at *λ* = *iω*.

#### Mean perturbation - phase

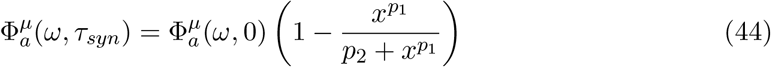

Where 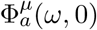 is the phase of the white noise response (36) when evaluated at *λ* = *iω*.

#### Variance perturbation - amplitude

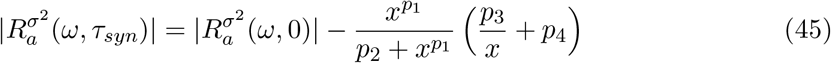

where 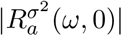 is the amplitude of the white noise response (36) when evaluated at *λ* = *iω*.

#### Variance perturbation - phase

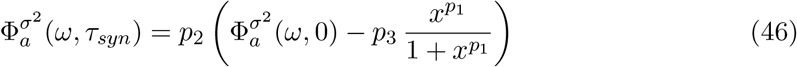

where 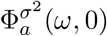 is the phase of the white noise response (36) when evaluated at *λ* = *iω*. We fitted the functions for *ω >* 0; responses for *ω <* 0 follow by even (amplitude) or odd (phase) symmetry. The best-fit parameters for the excitatory and inhibitory neuron are reported in Table 2.

**Table 2:** Fitted parameter values for response functions of E and I neurons.

| Parameter set | E neuron | I neuron |
| --- | --- | --- |
| Mean amplitude params | $p_1 = 1.807$ | $p_1 = 2.0571$ |
| | $p_2 = 2.1096$ | $p_2 = 4.2311$ |
| | $p_3 = -0.2989$ | $p_3 = -0.5295$ |
| Mean phase params | $p_1 = 1.0970$ | $p_1 = 1.1220$ |
| | $p_2 = 1.9558$ | $p_2 = 1.5426$ |
| Noise amplitude params | $p_1 = 2.3585$ | $p_1 = 2.6043$ |
| | $p_2 = 1.4783$ | $p_2 = 3.9899$ |
| | $p_3 = -0.5842$ | $p_3 = -0.9964$ |
| | $p_4 = 0.4065$ | $p_4 = 0.3834$ |
| Noise phase params | $p_1 = -1.8830$ | $p_1 = -2.7212$ |
| | $p_2 = 0.7653$ | $p_2 = 0.7712$ |
| | $p_3 = 0.3683$ | $p_3 = 0.2490$ |

Finally, we continued the fitted responses off the imaginary axis by evaluating the white-noise susceptibilities at the full complex *λ* and modulating their amplitude and phase with the fitted corrections evaluated at *ω* ≡ ℑ*λ*, thereby retaining the exact *λ*-dependence of the white-noise terms while using *ω*-only colored-noise corrections.

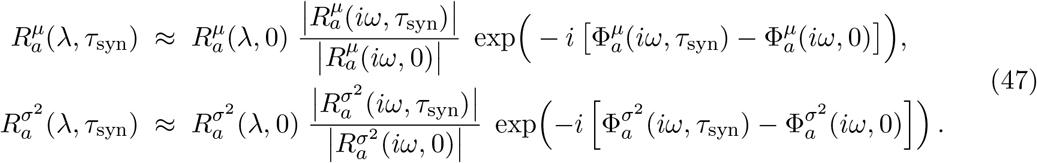

##### Oscillation frequency from rates

In Figs. 3b and 4b, we estimate the oscillation frequency of the population rates from simulations of 10 s. Let us define the population-averaged signal as *s*(*t*) = 0.8 *r_E_*(*t*) + 0.2 *r_I_*(*t*). We compute its (one-sided) power spectral density *S*(*ω*) via Welch’s method (Hamming window 1 s, 50% overlap), restricting to *ω* ∈ [5, 150] Hz. Let *ω*^⋆^ denote the frequency of the dominant spectral peak. To quantify uncertainty, we perform a jackknife over the *S* Welch segments: for each segment *s* = 1, …, *S*, we obtain a peak estimate 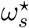, and report the segment mean and standard deviation,

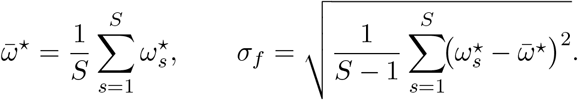

## 6 Supplementary Figures

**Figure S1:**
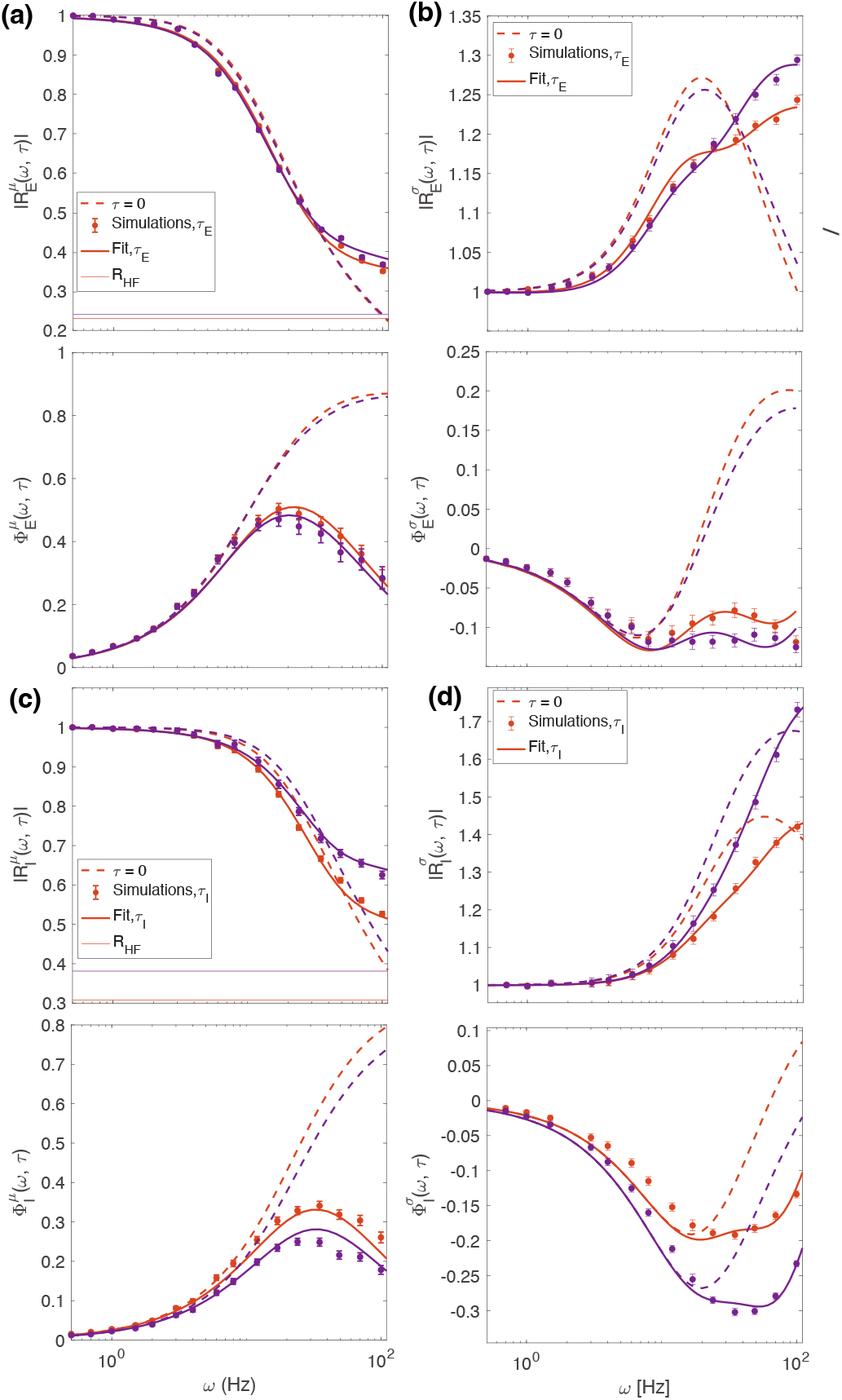
Response functions from mean field analysis. (a) Susceptibility to mean input current perturbations for E neurons. Top: Amplitude; Bottom: Phase. Dots represent simulations of uncoupled neurons receiving colored noise inputs, with statistics matching the full recurrent network at *g* = 3.6, *ν_ext_*/*ν_θ_* = 0.91 (orange); and *g* = 3.6, *ν_ext_*/*ν_θ_* = 1.3 (purple). Dotted lines show corresponding mean-field susceptibilities for zero synaptic time constants. Solid lines indicate the fit of a parametric mapping function transforming analytic white-noise susceptibilities to colored-noise counterparts (see Methods). Horizontal thin lines denote theoretical high-frequency limits (see eq. 39). (b) Susceptibility to input variance perturbations for E neurons. Details as in (a). (c–d) Susceptibilities for I neurons. Identical to (a) and (b), respectively, but calculated for the inhibitory population.

**Figure S2:**
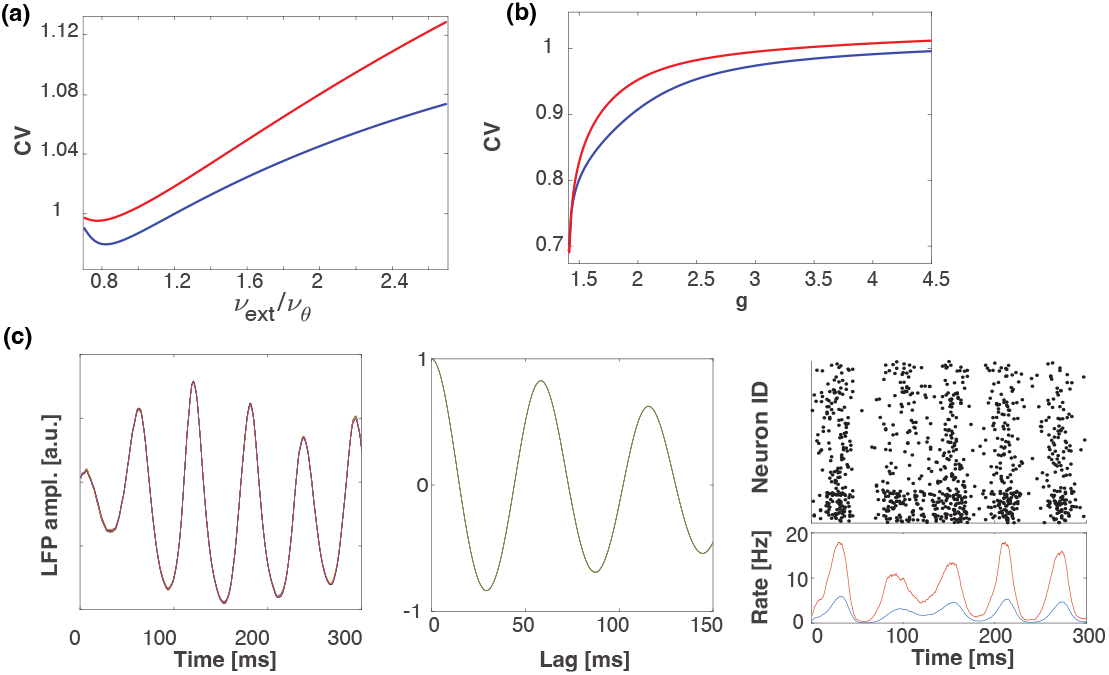
(a) Coefficient of variation (CV) of inter-spike intervals predicted from mean-field theory as a function of external input strength *ν*_ext_ at fixed inhibitory gain *g* = 3.65. This range corresponds to the simulations in Fig. 3c (bottom two rows). The CV remains near unity, indicating highly irregular single-neuron firing. CV as a function of the inhibitory gain *g* at fixed *ν*_ext_*/ν_θ_* = 1. At lower *g* values, where recurrent inhibition is weaker, the CV decreases slightly as spiking activity becomes more oscillatory, as illustrated in (c). (c) Representative network dynamics simulations at *g* = 2.5, *ν*_ext_*/ν_θ_* = 1. Left: LFP amplitude; middle: LFP autocorrelation function; right: raster plot of 800 excitatory (top) and 200 inhibitory (bottom) randomly selected neurons, overlaid with population firing rates for E (blue) and I (red) neurons.

**Figure S3:**
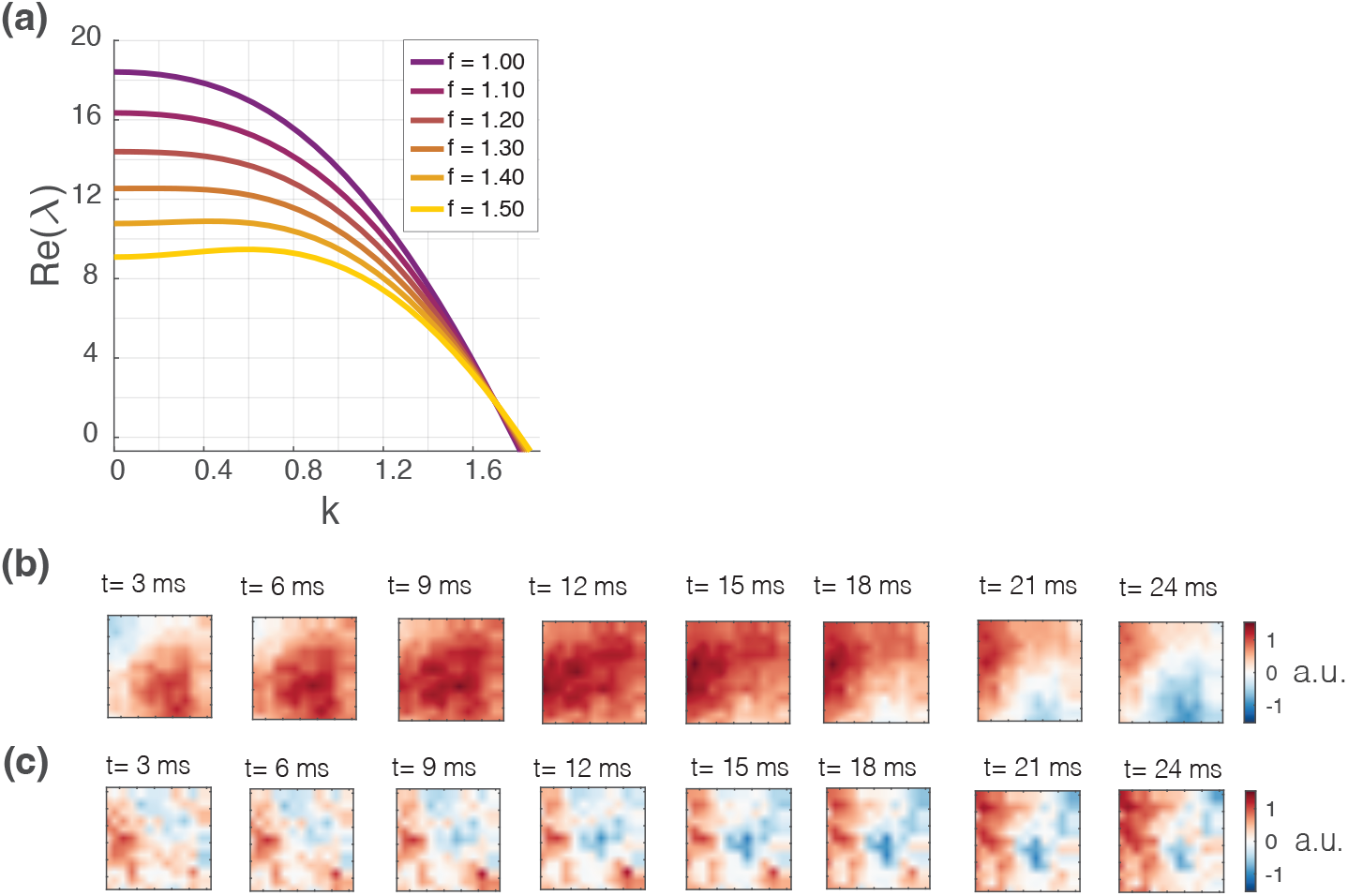
Model with homogeneous delays. (a) Real part of the largest eigenvalue, Re(*λ*), as a function of the wavenumber *k*. Colors indicate different delay magnitudes. Baseline delays are 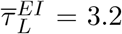 ms and 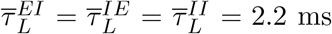, scaled by the factor *f* shown in the insert. Here, *g* = 3.625 and *ν*_ext_*/νθ* = 0.33. (b) Example of a wave event detected in simulations, for parameters within the spatio-temporal instability regime. (c) Example of activity when the homogeneous state is stable.

**Figure S4:**
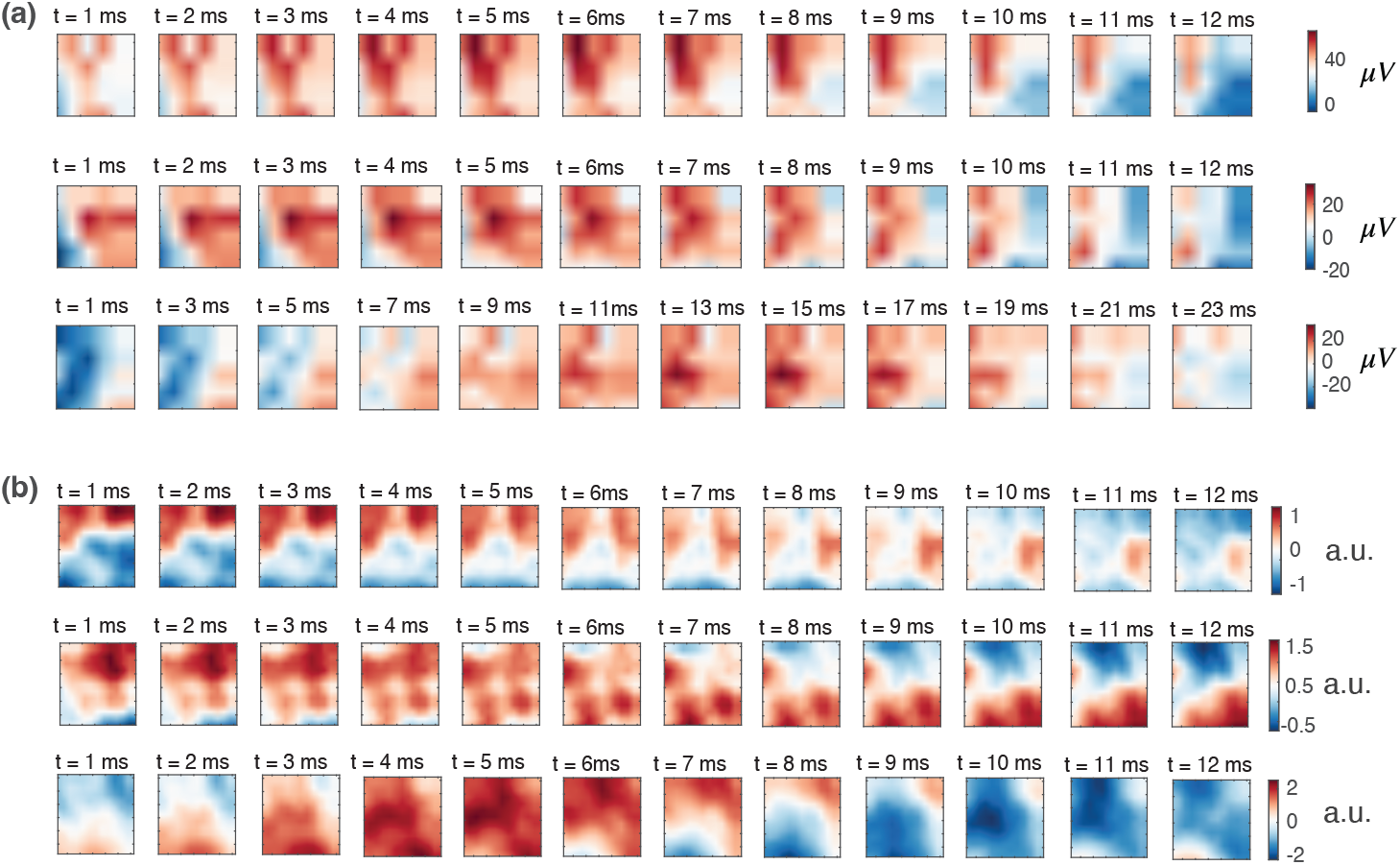
Spatiotemporal activity patterns: movement execution vs. asynchronous phase. (a) Example of LFP wave events detected from recordings during movement execution. For the purpose of comparison with simulations, the top row shows a section of the Utah array with 5×5 channels. (b) Example of wave events detected from simulations when the network is in the asynchronous phase, with virtual recording sites placed every 400 *µ*m.

**Figure S5:**
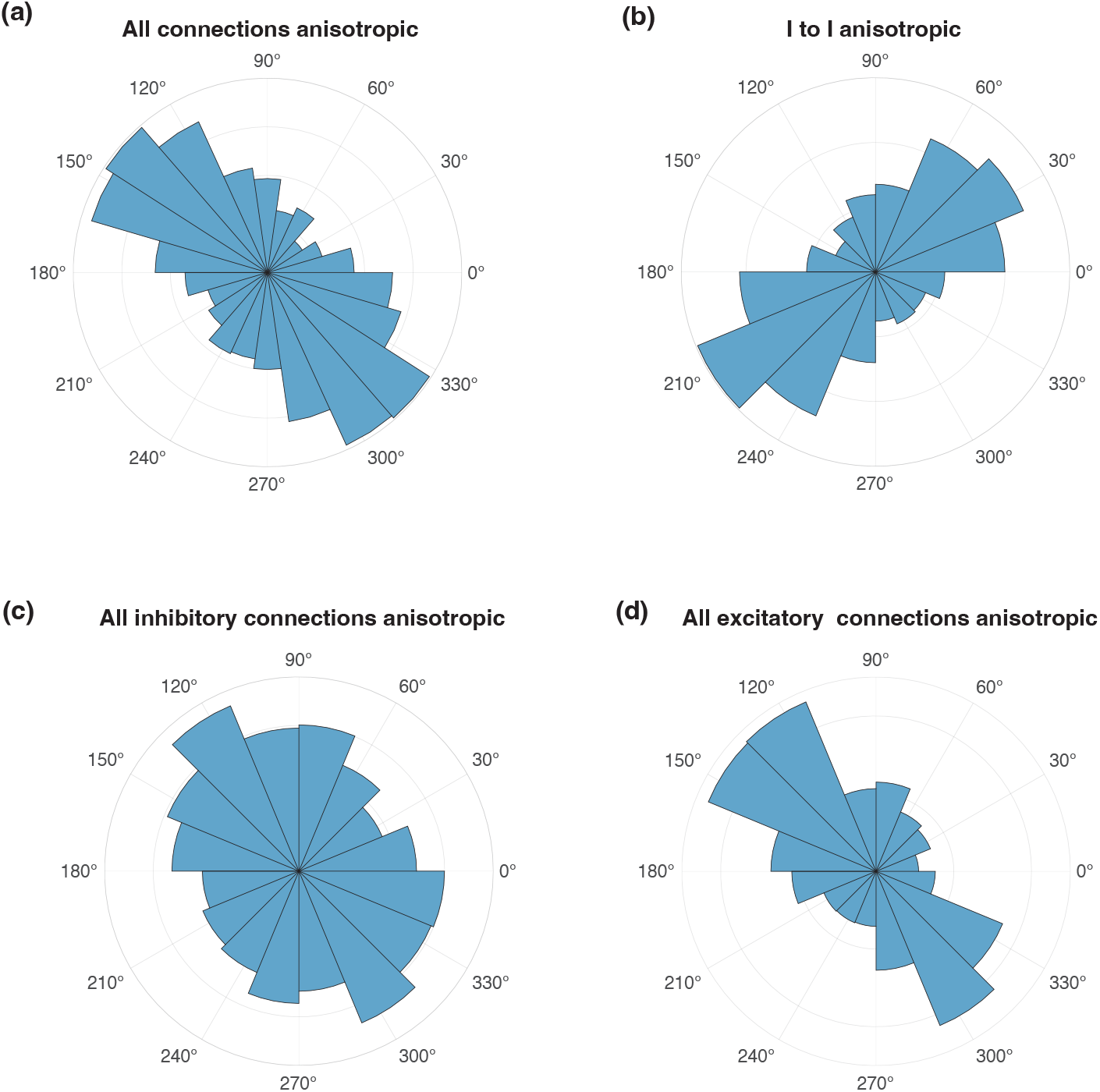
Effect of pathway-specific anisotropy on wave propagation direction. (a–d) Simulated directions of wave propagation for networks in which an anisotropy with fixed orientation of 45^◦^ relative to the *x*-axis is applied to different subsets of recurrent connections. For each condition, we show the distribution of propagation directions extracted from the phase gradients. (a) All recurrent connections are anisotropic (*ρ_EE_* = *ρ_IE_* = *ρ_EI_* = *ρ_II_* = 0.6). (b) Only I→I connections are anisotropic (*ρ_II_* = 0.6). (c) Only inhibitory connections are anisotropic (*ρ_EI_* = *ρ_II_* = 0.6). (d) Only excitatory connections are anisotropic (*ρ_EE_* = *ρ_IE_* = 0.6). In panels (a), (c), and (d), waves propagate predominantly along the axis of minimal spatial spread of the connectivity kernel, whereas in panel (b) they propagate along the axis of maximal spread. Results in all panels come from simulating 10 s of activity for 10 random realizations of the network.

**Figure S6:**
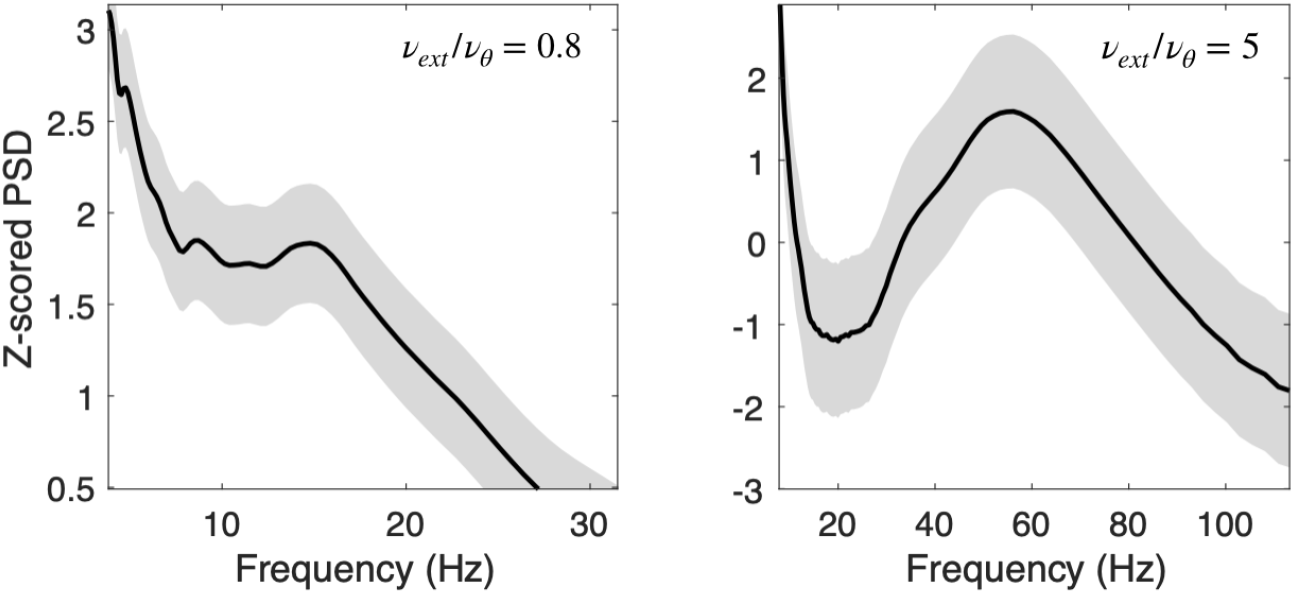
Normalized power spectral density from simulations, mean ± standard deviation over 3 trials. Left, external drive close to the lower boundary of the oscillatory regime. Right, external inputs that drive the network well above the asynchronous regime.

